# Cocaine Regulates Antiretroviral Therapy CNS Access Through Pregnane-X Receptor-Mediated Drug Transporter and Metabolizing Enzyme Modulation at the Blood Brain Barrier

**DOI:** 10.1101/2023.07.28.551042

**Authors:** Lisa B. Fridman, Stephen Knerler, Amira-Storm Price, Rodnie Colón Ortiz, Alicia Mercado, Hannah Wilkins, Bianca R. Flores, Benjamin C. Orsburn, Dionna W. Williams

## Abstract

**Background:** Appropriate interactions between antiretroviral therapies (ART) and drug transporters and metabolizing enzymes at the blood brain barrier (BBB) are critical to ensure adequate dosing of the brain to achieve HIV suppression. These proteins are modulated by demographic and lifestyle factors, including substance use. While understudied, illicit substances share drug transport and metabolism pathways with ART, increasing the potential for adverse drug:drug interactions. This is particularly important when considering the brain as it is relatively undertreated compared to peripheral organs and is vulnerable to substance use-mediated damage.

**Methods:** We used an *in vitro* model of the human BBB to determine the extravasation of three first-line ART drugs, emtricitabine (FTC), tenofovir (TFV), and dolutegravir (DTG), in the presence and absence of cocaine, which served as our illicit substance model. The impact of cocaine on BBB integrity and permeability, drug transporters, metabolizing enzymes, and their master transcriptional regulators were evaluated to determine the mechanisms by which substance use impacted ART central nervous system (CNS) availability.

**Results:** We determined that cocaine had a selective impact on ART extravasation, where it increased FTC’s ability to cross the BBB while decreasing TFV. DTG concentrations that passed the BBB were below quantifiable limits. Interestingly, the potent neuroinflammatory modulator, lipopolysaccharide, had no effect on ART transport, suggesting a specificity for cocaine. Unexpectedly, cocaine did not breach the BBB, as permeability to albumin and tight junction proteins and adhesion molecules remained unchanged. Rather, cocaine selectively decreased the pregnane-x receptor (PXR), but not constitutive androstane receptor (CAR). Consequently, drug transporter expression and activity decreased in endothelial cells of the BBB, including p-glycoprotein (P-gp), breast cancer resistance protein (BCRP), and multidrug resistance-associated protein 4 (MRP4). Further, cytochrome P450 3A4 (CYP3A4) enzymatic activity increased following cocaine treatment that coincided with decreased expression. Finally, cocaine modulated adenylate kinases are required to facilitate biotransformation of ART prodrugs to their phosphorylated, pharmacologically active counterparts.

**Conclusion:** Our findings indicate that additional considerations are needed in CNS HIV treatment strategies for people who use cocaine, as it may limit ART efficacy through regulation of drug transport and metabolizing pathways at the BBB.

## BACKGROUND

The current treatment strategy for the human immunodeficiency virus-1 (HIV) includes the use of antiretroviral therapies (ART) that target distinct steps of the viral life cycle, including entry, reverse transcription, and integration. ART administration occurs in a combinatorial fashion wherein two to three drugs from differing classes, termed combined ART (cART), are co-administered to facilitate suppressed viral replication. As a result, cART has revolutionized the HIV epidemic by decreasing the rates of opportunistic infections, acquired immunodeficiency syndrome (AIDS), HIV-related morbidity and mortality, and HIV transmission while increasing the lifespan for people living with the virus ^1–4^. However, treating HIV in tissues, including the brain, remains a challenge due to the limited potential of ART to bypass tissue barriers like the blood brain barrier (BBB) ^5–11^. This limited tissue access prevents organs from receiving adequate ART concentrations necessary for complete viral suppression and results in the development of sanctuaries that remain a major barrier to HIV eradication efforts ^12, 13^. Thus, it is imperative to understand more completely the mechanisms that impact ART access to the brain and other tissue sanctuaries.

It is not surprising that ART has difficulty traversing the BBB, as ∼98% of all therapeutic drugs face this same obstacle ^14, 15^. ART disposition in the brain requires navigating a complex and nuanced interplay between transporter proteins and drug metabolizing enzymes ^16^. Membrane-associated drug transporters facilitate influx into and efflux out of the CNS and primarily belong to ATP-binding cassette (ABC) and solute carrier (SLC) transporter superfamilies. While small, lipophilic, and uncharged molecules can passively diffuse through the BBB, large, hydrophobic, and charged molecules require facilitated and/or active transport by ABC and SLC transporters. A non-exhaustive list of the ABC and SLC transporters known to interact with ART include: P-glycoprotein (P-gp), multidrug resistance-associated protein 1 (MRP1), multidrug resistance-associated protein 2 (MRP2), multidrug resistance-associated protein 4 (MRP4), breast cancer resistance protein (BCRP), equilitative nucleoside transporter (ENT1), organic anion transporter 1 (OAT1), organic anion transporter 3 (OAT3), and organic anion-transporting polypeptide 1A2 (OATP1A2) ^17–19^. Drug metabolism is another obstacle that must be faced at the BBB, as phase I and II enzymes facilitate biotransformation through the addition of moieties (including oxygen and glucuronide groups) that increase hydrophilicity and aid in excretion. Of these, cytochrome P450 (CYP) 3A4 (CYP3A4) is among the most notable as it interacts with multiple ART classes, including reverse transcriptase inhibitors, protease inhibitors, entry inhibitors, and integrase inhibitors ^20–22^. Biotransformation is another aspect of drug metabolism particularly relevant for the ART’s administered as prodrugs, as it involves the addition of molecules to the parent drug required for becoming pharmacologically active. Adenylate kinases (AK) are one example of this, which, through their phosphorylation activity, facilitate the antiviral capacity of ART ^23–25^.

Drug transport and metabolism mechanisms are complex and encompass an interconnected web where simultaneous competing mechanisms occur due to wide and overlapping substrate specificity. For example, the reverse transcriptase inhibitor tenofovir (TFV) is the substrate of BCRP, MRP2, MRP4, OAT1, OAT3, and ENT1 while it can also induce P-gp, MRP1, MRP2, and MRP3 ^18, 19^. While not a substrate of CYP3A4, TFV can inhibit other CYP isoforms and requires kinase-mediated phosphorylation to become pharmacologically active ^24, 26–30^. It is also important to consider the intertwined nature of drug transport and metabolism mechanisms during HIV due to the combinatorial nature of cART. The co-administration of multiple drugs with overlapping specificity for drug transporters and metabolizing enzymes requires the interplay of multiple pathways to maintain appropriate ART concentrations and therapeutic efficacy. ART is not the only factor that must be taken into consideration, particularly as it pertains to treating sanctuary sites. One must consider the person behind the disease and the unique factors in their lives that may impact the ability of ART to work effectively, including diet, age, sex, and racial and/or ethnic background ^31–37^. For example, polypharmacy is highly prevalent in people living with HIV as many individuals also receive treatment for comorbid diseases ^38^. This creates the opportunity for drug:drug interactions that may change the pharmacologic profile of ART.

Substance use is another important, yet understudied, factor when evaluating ART access to the brain. Substance use is inextricably linked with the HIV epidemic and increases the risk of HIV acquisition ^39–42^. Further, the rate of substance use is higher among people living with HIV compared to seronegative individuals ^43–47^. Of importance, substance use is associated with poorer HIV outcomes, which is often attributed to decreased ART adherence ^48–52^. However, it is unlikely that every person with HIV who consumes illicit substances discontinues taking ART as prescribed. While this may certainly occur for some individuals, the molecular consequences of substance use on ART efficacy should also be considered. Interestingly, adverse drug:drug reactions exist between substances of abuse and ART, due to shared drug transport and metabolism pathways, which can lead to decreased ART efficacy, increased toxicity, and poorer outcomes for people living with HIV ^53–59^. Additionally, the impact of substance use on ART efficacy in the brain is of particular importance, as illicit substances are well known to impact BBB and CNS function ^60–67^.

Extensive regulatory mechanisms exist to ensure the proteins involved in drug transport and metabolism fulfill their endogenous responsibilities, while also promoting detoxification of the cell and xenobiotic clearance. Two players are tasked with being the master orchestrators of these pathways: the nuclear receptors pregnane-X receptor (PXR) and constitutive androstane receptor (CAR) ^68^. These ligand-activated transcription factors are xenobiotic sensors that, following activation, coordinately regulate genes encoding drug transporters and drug metabolizing proteins ^69–71^. They have overlapping activity and interact widely with licit and illicit pharmacologically active substances ^72–74^. PXR and CAR are highly expressed at the BBB where they regulate the activity of drug transporter and metabolizing enzymes, including P-gp, BCRP, and MRP2 ^75–82^. Additionally, their downregulation can decrease the expression and activity of drug transporters. Of importance, ART and substances of abuse are capable of inducing changes in drug transporter and metabolizing enzyme expression through interactions with PXR and CAR, which may alter their anti-HIV pharmacokinetic properties ^18, 19, 83, 84^. Moreover, the modulation of PXR and CAR may promote adverse drug-drug interactions between substances of abuse and ART that can result in decreased antiviral efficacy and potentially treatment failure – especially in the brain and other tissue reservoirs.

We used cocaine as a model illicit substance to evaluate the impact of substance use on ART CNS availability through interactions at the BBB. Our study centered on three ART drugs that represent a first-line HIV regimen: emtricitabine (FTC), TFV, and dolutegravir (DTG). Using a transwell model of the human BBB, we evaluated the ability of FTC, TFV, and DTG to cross from the apical to basolateral chamber in the presence and absence of cocaine, or lipopolysaccharide (LPS) as a control. We identified an inherent differential capacity of ART to cross the BBB, where TFV had the highest extravasation rate followed by FTC. DTG’s migration across the BBB was below quantifiable limits. Cocaine, but not LPS, altered ART’s ability to cross the BBB where it increased FTC, but decreased TFV. Unexpectedly, cocaine’s effects on ART extravasation did not cause BBB disruption. Instead, cocaine decreased PXR that resulted in altered drug transporter and drug metabolizing enzyme expression and activity. Of note, cocaine’s effects were specific to PXR as CAR remained unchanged. Our findings demonstrate that cocaine can regulate ART bioavailability and efficacy in the CNS by regulating drug transport and metabolism activity at the BBB. Further, our study suggests substance use must be taken into consideration in ART prescription recommendations to ensure all people with HIV have an equal chance to achieve viral suppression, especially in sanctuaries like the brain.

## METHODS

### Cells

Primary human astrocytes (ScienCell Research Laboratories, Carlsbad, CA) were grown to confluence in Basal Medium Eagle (Thermo Fisher Scientific, Waltham, MA) buffered to pH ranging from 7.2-7.5 with 2.2 g/L sodium bicarbonate and 15 mM HEPES (Gibco, Grand Island, New York). Media was supplemented with 2% fetal bovine serum (FBS) (R&D Systems, Minneapolis, MN), 1% penicillin-streptomycin 10,000U/mL (Gibco), and 1% astrocyte growth supplement (ScienCell Research Laboratories). Astrocytes were used at passages 3-4 for all experiments.

Primary human brain microvascular endothelial cells (Cell Systems, Kirkland, WA) were grown to confluence on tissue culture plates coated with 0.2% gelatin (Thermo Fisher Scientific) in medium 199 (M199) (Gibco) buffered to pH ranging from 7.2-7.5 with 2.2 g/L sodium bicarbonate and 15 mM HEPES (Gibco). Complete M199 media (M199C) was comprised of 20% heat-inactivated newborn calf serum (Gibco), 1% penicillin-streptomycin 10,000U/mL (Gibco), 25 mg/L heparin (Sigma, St. Louis, MO), 5% heat-inactivated human serum AB (GeminiBio, Sacramento, CA), 50 mg/L ascorbic acid (Sigma), 7.5 mg/L endothelial cell growth supplement (Sigma), 2 mM L-glutamine (Gibco), and 5 mg/L bovine brain extract (Lonza, San Diego, CA). Endothelial cells were used at passages 9-16 for all experiments.

### *In vitro* Model of the Human BBB

Our *in vitro* transwell model of the human BBB model was made as previously described ^85–91^. Briefly, astrocytes (1×10^5^ cells/insert) were seeded on the underside of a tissue culture insert comprised of a polycarbonate membrane with 3 μM pores (Falcon, Corning, NY) and allowed to adhere for four hours at 37°C, 5% CO_2_ while continually being fed in 5-30 minute intervals with M199C. The tissue culture inserts were then inverted and transferred to a 24-well tissue culture plate (Falcon) containing M199C. Endothelial cells (4×10^4^ cells/insert) were seeded into the upperside of the insert that was pre-coated with 0.2% gelatin (Thermo Fisher Scientific). The cells grew to confluence at 37°C, 5% CO_2_ over three days, during which time the astrocyte processes penetrated through the pores to establish contact with the endothelial cells and seal the barrier. The BBB model was used for experiments following four days of culture.

### Evans Blue Albumin Permeability Assay

Permeability of our *in vitro* BBB model was evaluated using Evans Blue dye conjugated to albumin (EBA). To prepare the EBA dye, 0.45% Evans Blue (Sigma) was conjugated to bovine serum albumin (Thermo Fisher Scientific) by incubation at 37°C overnight while rotating continuously. Excess unbound dye was removed by ice cold ethanol washes that consisted of precipitation with 100% molecular grade ethanol (The Warner Graham Company, Cockeysville, MD) at -80°C for 30 minutes, centrifugation at 4°C at maximum speed (21,130 g) for 10 minutes, removal of the unbound dye-containing supernatant, mechanical dissociation of the albumin pellet, and washing with ice cold ethanol prior to repeating albumin precipitation at -80°C. The ethanol precipitation and washes were repeated for ∼40 cycles until the supernatants were clear and all unbound Evans Blue dye was removed.

BBB permeability was determined by adding EBA to the apical portion of our transwell model for 30 minutes at 37°C, 5% CO_2_ and allowing it to pass into the basolateral chamber containing phenol red free Dulbecco’s Modified Eagle Medium (Gibco). After the indicated time, the media contained in the basolateral portion was collected and the absorbance spectrophotometrically evaluated at 620 nm. EBA dye added to phenol red free Dulbecco’s Modified Eagle Medium served as a positive control to determine quantitation of the absorbance value corresponding to a complete breach of the BBB.

### ART BBB Extravasation Assay

FTC, TFV, and DTG (all from Toronto Research Chemicals, Toronto, Canada) were reconstituted to 10 μM in M199C and added the apical portion of the BBB model in the presence or absence of 10 ng/mL LPS (Sigma) or 10 μM cocaine hydrochloride (NIDA Drug Supply Program, Research Triangle Park, NC) for 24 hours at 37°C, 5% CO_2_. M199C alone was used as a negative control. After the indicated time, the media contained in the basolateral portion was collected, aliquoted, and stored at -80°C until quantitation by tandem liquid chromatography-mass spectrometry analyses. There were no freeze/thaw cycles before quantitation.

### ART Concentration Determination

The concentrations of FTC, TFV, and DTG that passed through the BBB model were determined using validated liquid chromatographic-mass spectrometric methods by the Clinical Pharmacology Analytic Laboratory at the Johns Hopkins University School of Medicine, as previously described ^92, 93^. Briefly, FTC and TFV were analyzed in positive mode using a TSQ Vantage^®^ triple quadrupole mass spectrometer coupled with a HESI II^®^ probe (Thermo Scientific). The analytical run time was 8 min, and the assay lower limits of quantitation were 5 and 1 ng/mL for FTC and TFV, respectively. DTG was analyzed using an API 5000 mass analyzer (SCIEX, Redwood City, CA, USA) interfaced with a Waters Acquity UPLC system (Waters Corporation, Milford, MA, USA). The analytical run time was 2-5 minutes, and the assay lower limit of quantitation was 100 ng/mL. DTG concentrations below 100 ng/mL were reported as below the limit of quantitation. Assays were validated in accordance with the FDA Guidance for Industry, Bioanalytical Method Validation recommendations and by the Clinical Pharmacology Quality Assurance program ^94, 95^. All performance parameters were acceptable.

### Endothelial Cell Cocaine Treatment

When 80% confluent, primary human endothelial cells were treated with 0.01-100 μM cocaine hydrochloride for 24 hours, after which time they were used in all subsequent downstream assays. Treatment with vehicle was used as a control.

### Quantitative RT-PCR

Endothelial cells were lysed with Buffer RLT Plus (Qiagen, Germantown, MD) supplemented with 1% β-mercaptoethanol (Sigma). Total RNA was isolated using the RNeasy Mini Kit (Qiagen) according to the manufacturer’s protocol with the modification of on column DNase digestion using RQ1 RNase free DNase (Promega, Madison, WI) in the enzyme mix. Complementary DNA (cDNA) synthesis was performed using 1 μg of total RNA with the iScript cDNA Synthesis Kit (Bio-Rad, Hercules, CA). The genes encoding for the zonula occludens-1 (Zo-1), OAT1, OAT3, ENT1, OATP1A2, OATP2A1, BCRP, P-gp, MRP1, MRP4, MRP5, CYP3A4, PXR, CAR, and 18S proteins were evaluated by qRT-PCR using a Taqman Gene Expression Assay (Thermo Fisher Scientific, Waltham, MA) using the BioRad CFX96 Real-Time System with cycling conditions optimized for the TaqMan Fast Advanced Master Mix (enzyme activation at 95°C for 20 seconds, 40 cycles of denaturing at 95°C for 1 second, and annealing/extending at 60°C 20 seconds). Results were normalized to 18S and presented as a fold change relative to the treatment vehicle using the 2-^ΔΔCt^ method, where the vehicle treated group was set to 1.

### Western Blot

Endothelial cells were lysed with 1X RIPA buffer (Cell Signaling Technology, Danvers, MA) supplemented with 1X protease/phosphatase inhibitor (Cell Signaling Technology). Total protein concentrations were determined by Bradford Assay with the Bio-Rad Protein Assay Dye reagent concentrate (Bio-Rad) following the manufacturer’s instructions. Forty μg of protein was electrophoresed on a 4-12% polyacrylamide gel (Bio-Rad) and transferred to nitrocellulose membranes (Amersham Biosciences, Woburn, MA). Membranes were blocked for two hours at room temperature with 5% nonfat dry milk (Lab Scientific bioKEMIX Inc., Danvers, MA) and 3% bovine serum albumin (Thermo Fisher Scientific) in 1X Tris-Buffered Saline (Quality Biological, Gaithersburg, MD) containing 0.1% Tween-20 (TBS-T, Sigma). Blots were probed with antibodies with specificity to Zo-1, claudin-5, occludin, OAT1, OAT3, ENT1, OATP1A2, OATP2A1, BCRP, P-gp, MRP1, MRP4, MRP5, CYP3A4, PXR, CAR, AK1, AK2, AK5 or AK6, overnight at 4°C, washed with TBS-T, and probed with the appropriate secondary antibody for one hour at room temperature. Antibody details are provided in **Table 1**. All antibodies were titered to determine optimal concentrations. Western Lightning Plus-ECL (PerkinElmer, Waltham, MA) was used as chemiluminescence substrate and the signal detected with the Azure Biosystems c600 Imager (Azure Biosystems, Dublin, CA). As a loading control, membranes were stripped with Restore Plus Western Blot Stripping Buffer (Thermo Fisher Scientific) and reprobed with antibody against β-Actin HRP for one hour at room temperature. Densitometric analysis was performed using ImageJ (Version 1.53t, NIH, Bethesda, MD) to quantitate the band density (pixels, arbitrary units) for all evaluated proteins. Relative band intensity for each protein of interest was determined by calculating its pixel ratio with β-actin. The vehicle treated group was set to 1 and the relative fold change in protein expression relative to vehicle determined.

**Table 1.** Details regarding antibody name, catalogue number, clone, concentration, and company from which it was obtained are provided for antibodies used in the study.

| Antibody | Company | Catalogue Number | Clone | Concentration for Immunofluorescence | Concentration for Western Blot |
| --- | --- | --- | --- | --- | --- |
| GFAP(AF488) | Invitrogen | 53-9892-82 | GA5 | 10 µg/mL |  |
| GLUT-1 | Invitrogen | PA5-16793 | Poly | 1:100 |  |
| CD71 | Invitrogen | 14-0719-82 | OKT9 | 5 µg/mL |  |
| PECAM(FITC) | Invitrogen | 11-0311-82 | 390 | 5 µg/mL |  |
| VE-cadherin | Invitrogen | 14-1449-82 | 16B1 | 5 µg/mL |  |
| Zo-1 | Invitrogen | 61-7300 | Poly | 10 µg/mL | 1 µg/mL |
| Occludin | Invitrogen | 71-1500 | Poly | 4 µg/mL | 0.5 µg/mL |
| Claudin 5 | Invitrogen | 35-2500 | 4C3C2 | 1:25 | 1:1000 |
| OAT1 | Invitrogen | PA5-26244 | Poly | 1:100 | 1:1000 |
| OAT3 | Invitrogen | PA5-76143 | Poly | 1:100 | 1:1000 |
| ENT1 | Proteintech | 11337-1-AP | Poly | 1:100 |  |
| ENT1 | Invitrogen | PA5-116461 | Poly |  | 1:500 |
| OATP1A2 | Invitrogen | PA5-42445 | Poly | 1:100 | 1:1000 |
| OATP2A1 | Invitrogen | PA5-98789 | Poly | 1:200 | 1:1000 |
| BCRP | Novus | NBP2-22124 | 3G8 | 1:200 | 1:2000 |
| P-gp | Invitrogen | PA5-61300 | Poly | 2 µg/mL |  |
| P-gp | Abcam | ab261736 | Poly |  | 0.5 µg/mL |
| MRP1 | Abcam | ab24102 | MRPm 5 | 1:50 | 1:200 |
| MRP4 | Cell Signaling | 12705 | D2Q2O | 1:200 | 1:1000 |
| MRP5 | Invitrogen | PA5-18965 | Poly | 10 µg/mL |  |
| MRP5 | Abcam | Ab180724 | Poly |  | 1:1000 |
| CYP3A4 | Invitrogen | MA5-17064 | 3H8 | 1:200 | 1:1000 |
| PXR | Invitrogen | PA5-72551 | Poly | 1:50 | 1:1000 |
| CAR | R&D Systems | PP-N4111-00 | N4111 | 10 µg/mL | 2 µg/mL |
| AK1 | Invitrogen | PA5-52297 | Poly | 1 µg/mL | 0.1 µg/mL |
| AK2 | Proteintech | 11014-1-AP | Poly | 1:200 | 1:1000 |
| AK5 | Invitrogen | PA5-53933 | Poly | 1.2 µg/mL | 0.09 µg/mL |
| AK6 | Novus | NBP2-67116 | B1-F4 | 1:100 | 1:2000 |
| B-Actin (HRP) | Cell Signaling | 12262 | 8H10D 10 |  | 1:5000 |
| Anti-rabbit (HRP) | Abcam | ab97051 | Poly |  | 1:2000 |
| Anti-mouse (HRP) | Abcam | ab97023 | Poly |  | 1:2000 |
| Anti-rabbit (AF488) | Invitrogen | A-11008 | Poly | 1 µg/mL |  |
| Anti-mouse (AF488) | Invitrogen | A-11001 | Poly | 0.67 µg/mL |  |
| Anti-goat (AF488) | Invitrogen | A-11078 | Poly | 1 µg/mL |  |

### Proteomics

Endothelial cells were treated with cocaine (10 µM) or vehicle for 24 hours at 37°C, 5% CO_2_, after which time whole cell proteins were extracted using a 5% sodium dodecyl sulfate and 50 mM triethylammonium bicarbonate lysis buffer. Protein concentration was determined using the Pierce BCA Protein Assay Kit (Thermo Fisher Scientific) following the manufacturer’s instructions. Proteins were digested into peptides using the S-Trap^TM^ 96-well plate (ProtiFi, Fairport, NY) following the manufacturer’s instructions. In brief, 100 μg of protein from each sample was solubilized in 5% SDS, reduced with 120 mM tris(2-carboxyethyl)phosphine (ProtiFi), alkylated using 500 mM methyl methanethiosulfonate (ProtiFi), acidified with 1.2% phosphoric acid (ProtiFi), trapped on column, and then digested by 10 μg MS-grade trypsin (Thermo Scientific). Once peptides were eluted, they were dried under vacuum centrifugation (Eppendorf, Enfield CT) overnight, resuspended in 100 μl 0.1% formic acid in H_2_O (Thermo Fisher Scientific), and quantified using the Pierce Quantitative Colorimetric Peptide Assay kit (Thermo Scientific). Samples were diluted to 100 ng/μl and 2 μl were injected by an EasyNLC 1200 (Thermo Fisher Scientific) nanoflow liquid chromatography system coupled to a timsTOF FleX mass spectrometer (Bruker, Billerica, MA). Mobile phase A was 0.1% formic acid in H_2_O (Thermo Fisher Scientific) and mobile phase B was 0.1% formic acid (Thermo Fisher Scientific) in 80% acetonitrile/20% H_2_O (Thermo Fisher Scientific). Peptides passed through an Acclaim PepMap C18 100Å, 3 μm, 75 μm × 2 cm trap column (Thermo Scientific) followed by separation on a PepSep C18 100Å, 1.5 μm, 75 μm × 15 cm (Bruker) at a flow rate of 200 nl/min using the following 1 hr gradient: 10% - 35% B from 0-47 min, 35% - 100% B from 47–55 min, 100% B from 55 min-57 min, 100% - 5% B from 57 min-58 min, and 5% B from 58 min-60 min. The trap column equilibration used a 9 μl at 3.0 μl/min flow rate and the separation column equilibration used a 12 μl at 3.0 μl/min flow rate. Additionally, 1 wash cycle of 20 μl and a flush volume of 100 μl were used. Peptides were ionized using the CaptiveSpray source, with a capillary voltage of 1500V, dry gas flow of 3 l/min and temperature 180°C. Data were acquired using a positive ion mode diaPASEF method with a mass range from 100-1700 m/z and 1/K_0_ from 0.80 V_s_/cm^2^ to 1.35 V_s_/cm^2^ with 100 ms ramp time and 2 ms accumulation time. General tune parameters were: Funnel 1 RF = 300 Vpp, idCID Energy = 0 eV, Deflection Delta = 70 V, Funnel 2 RF = 200 Vpp, Multipole RF = 500 Vpp, Ion Energy = 5 eV, Low Mass = 200 m/z, Collision Energy = 10 eV, Collision RF = 1500 Vpp, Transfer Time = 60 μs, Pre Pulse Storage = 12 μs, and Stepping turned off. Tims tune parameters were: D1 = -20 V, D2 = -160 V, D3 = 110 V, D4 = 110 V, D5 = 0 V, D6 = 55 V, Funnel1 RF = 475 Vpp, and Collision Cell In = 300V. Resulting spectra were uploaded to Spectronaut 17.1 (Biognosys, Cambridge, MA). Peptides were identified and quantified using the directDIA analysis default settings with the proteotypicity filter set to “Only Proteotypic” and a variable modification set to “methylthio.” MS1 protein group quantifications and associated protein group UniProt numbers and molecular weights were exported from Spectronaut. MS1 protein group quantifications, UniProt numbers, and molecular weights were imported into Perseus for use of the proteomic ruler plug-in, as previously described ^96^. The default proteomic ruler plug-in settings were used with histone proteomic ruler as the scaling mode, ploidy set to two, and total cellular protein concentration set to 200 g/l. Protein concentrations (nM) and copy numbers as estimated by the proteomic ruler were used f or analysis.

### Flow Cytometry

Endothelial cells were gently recovered from tissue culture plates using TrypLE Express (Invitrogen, Grand Island, NY) to maintain surface antigen expression for 10-15 minutes at 37°C, 5% CO_2_ as previously described ^91^. After recovery, the cells were washed once with phosphate buffered saline (PBS, Gibco) and extracellular immunostaining performed. The cells were washed once with cold flow cytometry buffer (PBS supplemented with 2% Human Serum) (Corning, Manassas, VA) and 500,000 cells per tube were stained with fluorochrome-coupled antibodies specific for CD54/ICAM, CD31/PECAM-1, F11r/JAM-A, CD166/ALCAM, or corresponding isotype-matched negative control antibodies (BD Biosciences, Franklin Lakes, NJ) in the dark, on ice for 30 minutes. Antibody details are listed in **Table 1**. All antibodies were titered to determine optimal concentrations for staining. Following staining, the cells were washed with cold flow cytometry buffer and fixed with 2% paraformaldehyde (Electron Microscopy Sciences, Hatfield, PA). The samples were stored at 4°C wrapped in foil up to 1 week prior to flow cytometric analysis. Samples were filtered using BD FACS tubes with cell strainer caps with 35-µm pores (BD Biosciences) immediately before flow cytometric acquisition. At least 10,000 live, singlet events were acquired with a BD LSRFortessa cytometer and Diva software version 9 on the Windows 10 platform (BD Biosciences). Flow cytometric data were analyzed using FlowJo version 10.9 (FlowJo, Ashland, OR).

### Immunofluorescent Microscopy

Endothelial cells were seeded (1.6×10^4^ cells/dish) on 35 mm ibiTreat dishes (Ibidi USA, Madison, WI) coated with 0.2% gelatin, grown to 80% confluence, and treated with cocaine hydrochloride. Following treatment, cells were fixed with 4% paraformaldehyde (Electron Microscopy Sciences) for 15 minutes. The cells were stained with wheat germ agglutinin conjugated to Texas red (Thermo Fisher Scientific) for 10 minutes to facilitate identification of cell morphology through staining of the plasma membrane. Cells were then permeabilized in 0.01% Triton X-100 (Sigma) for one minute then blocked for two hours at room temperature in Dulbecco’s phosphate-buffered saline without calcium or magnesium (DPBS) (Thermo Fisher Scientific) containing 5 mM ethylenediaminetetraacetic acid (EDTA) (Sigma), 1% fish gelatin (Sigma), 1% essentially immunoglobulin-free bovine serum albumin (Sigma), 1% heat-inactivated human serum AB (GeminiBio, Sacramento, CA), and 1% goat serum (Vector Laboratories, Newark, CA). Cells were probed with antibodies with specificity to Zo-1, OAT1, OAT3, ENT1, OATP1A2, OATP2A1, BCRP, P-gp, MRP1, MRP4, MRP5, CYP3A4, PXR, CAR, AK1, AK2, AK5 or AK6, overnight at 4°C, washed three times with DPBS at room temperature, and probed with the appropriate Alexa Fluor 488 conjugated secondary antibody for one hour at room temperature. Isotype-matched controls, staining with only secondary antibodies, and unstained cells were used as negative controls and to account for autofluorescence and nonspecific signal. Antibody details are listed in **Table 1**. Cells were mounted with Ibidi mounting medium containing DAPI as a counterstain to identify nuclei (Ibidi USA, Madison, WI). Cells were imaged by fluorescent microscopy using the ECHO Revolution (San Diego, CA).

For evaluation of the BBB model, a surgical blade was used to outline the polycarbonate membrane from the tissue culture insert from the basolateral side, leaving only a small portion adhered to the insert. The polycarbonate membrane was carefully removed from the tissue culture insert, gently washed with PBS, and fixed with 4% paraformaldehyde (Electron Microscopy Sciences) for 20 minutes at 37°C, 5% CO_2_. The polycarbonate membranes were placed into an eight-well chamber slide (Ibidi USA), paying attention to place the apical or basolateral side facing downwards to stain the endothelial cells or astrocytes, respectively. The polycarbonate membranes were stained with wheat germ agglutinin conjugated to Texas red (Thermo Fisher Scientific) for 10 minutes to facilitate identification of cell morphology through staining of the plasma membrane. Cells on the polycarbonate membranes were then permeabilized in 0.01% Triton X-100 (Sigma) for one minute then blocked for two hours at room temperature in DPBS containing 5 mM EDTA (Sigma), 1% fish gelatin (Sigma), 1% essentially immunoglobulin-free bovine serum albumin (Sigma), 1% heat-inactivated human serum AB (GeminiBio, Sacramento, CA), and 1% goat serum (Vector Laboratories, Newark, CA). The polycarbonate membranes were probed with antibodies with specificity to GFAP or VE-Cadherin, overnight at 4°C, washed three times with DPBS at room temperature, and probed with the appropriate Alexa Fluor 488 conjugated secondary antibody for one hour at room temperature. Antibody details are listed in **Table 1**. All antibodies were titered to determine optimal concentrations. Isotype-matched controls, staining with only secondary antibodies, and unstained membranes were used as negative controls and to account for autofluorescence of the polycarbonate membranes and nonspecific signal. The polycarbonate membranes were stained with Ibidi mounting medium containing DAPI as a counterstain to identify nuclei (Ibidi USA, Madison, WI). The polycarbonate membranes were imaged by fluorescent microscopy using the ECHO Revolution (San Diego, CA).

Images were acquired 1-15 days post fixation using the ECHO Revolution in the inverted mode with a 20X Plan Apo objective (with a 1.4 numerical aperture) where three channels were used: blue to identify nuclei, red to identify cell morphology, and green to identify the protein of interest. The appropriate focal plane along the Z-axis was determining manually prior to image acquisition. The signal to noise ratio was maximized by optimizing the appropriate exposure time and incident light intensity prior to image acquisition to prevent saturation and minimize background. Identical acquisition settings, including exposure time and intensity, were used for all treatment conditions where images were acquired on the same day to minimize batch effects due to fluorescent fading/quenching. Twenty images were acquired for all treatment conditions, with the exception of Zo-1 for which 10 images were taken and the BBB inserts where 3-5 images were taken. (FIJI is Just) ImageJ v1.54b (National Institutes of Health) was used for image quantification.

### Efflux Transporter Activity Assay

Efflux transporter activity was determined by cellular efflux of rhodamine 123 (10 µM, Thermo Fisher Scientific), Hoechst 33342 (5 µg/mL, Thermo Fisher Scientific), and monobromobimane (10 µM, Thermo Fisher Scientific), fluorescent substrates with specificity for P-gp, BCRP, and MRP4, respectively. Endothelial cells were incubated with fluorescent substrate for 1 hour at 37°C, 5% CO_2_ to allow uptake into the cell, after which time fresh media added and the substrates allowed to efflux from the cells for four hours at 37°C, 5% CO_2_. To evaluate the impact of cocaine on efflux transporter activity, endothelial cells were pre-treated with 10 µM cocaine hydrochloride for 24 hours prior to addition of the fluorescent substrates. To evaluate the effect of PXR inhibition on efflux transporter activity, the endothelial cells were pre-treated with 10 µM of the PXR-specific inhibitor resveratrol (Sigma) 24 hours prior to addition of the fluorescent substrates. As a control, 10 µM ritonavir (Sigma), fumitremorgin C (Sigma), and ceefourin 1 (Tocris Bioscience, Minneapolis, MN) were added concomitantly with the media change and served as known inhibitors of P-gp, BCRP, and MRP4, respectively. Vehicle treatment was used as a negative control. Following treatments, cells were washed with PBS, detached from tissue culture plates with 0.5% Trypsin-EDTA (Gibco), and the cells washed with PBS again. The cells were filtered using BD FACS tubes with cell strainer caps with 35-µm pores (BD Biosciences) and immediately subject to flow cytometric acquisition where at least 10,000 singlet events were acquired with a BD LSRFortessa cytometer and Diva software version 9 on the Windows 10 platform (BD Biosciences). Flow cytometric data were analyzed using FlowJo version 10.9 (FlowJo, Ashland, OR).

### CYP3A4 Metabolic Activity Assay

Endothelial cells were plated in 96-black microplate with clear flat bottom (Corning, NY) coated with 0.2% gelatin (Thermo Fisher Scientific) at a density of 5,000 cells per well and cultured overnight at 37°C, 5% CO_2_ in M199C. After overnight culture, endothelial cells were pre-treated with cocaine (10 μM), positive control rifampicin (1 µM, Sigma) the PXR-specific inhibitor resveratrol (10 µM, Sigma), or vehicle control. Twenty-four hours post treatment, the cells were washed twice with PBS and incubated with 2 μM of the CYP3A4 fluorogenic probe substrate, 7-benzyloxy-4-trifluoromethylcoumarin (BFC, Sigma), for 90 minutes at 37°C, 5% to permit its oxidative enzymatic conversion to the fluorescent metabolite 7-hydroxy-4-trifluoromethylcoumarin (HFC). After this period, fluorometric quantitation was performed using the Spectra Max iD5 (Molecular Devices, San Jose, CA) microplate reader at excitation and emission wavelengths of 405/535 nm.

Baseline subtraction was performed by subtracting the RFU at the initial timepoint (0) for vehicle treatment from all other conditions. The ratio between RFU and time (minutes) was taken to calculate CYP3A4 velocity (RFU/minutes). Only the linear portion of the curve (0-20 minutes) was used for analysis.

### Statistical Analysis

Three independent experiments comprised of four technical replicates were performed for ART BBB extravasation assays. Seven independent experiments comprised of three technical replicates were performed for BBB permeability assays. Samples for qRT-PCR’s were run in triplicate. Liquid chromatography/mass spectrometry and proteomic assays were run with three independent sample injections. All remaining *in vitro* experiments were repeated in at least n≥5 independent experiments.

Raw files without compression were used for immunofluorescent microscopy quantification where 250-500 cells were analyzed for each treatment, with the exception of Zo-1 where 70-100 cells were used. Three regions of interest (ROI) were used to facilitate quantitation of fluorescent signal: background, nuclei, and cells. The background signal was determined using the red channel where rectangular ROI’s were drawn in regions containing no cells. Background ROI’s were superimposed onto the green channel for the protein of interest and the average intensity was measured. Nuclei were segmented from the background using the blue channel and creating a binary image using Otsu thresholding. Nuclear ROI’s were created using the binary image and the particle analyzer in FIJI (size >1000-pixel units, circularity between 0.00-1.00). Nuclear ROI’s were superimposed onto the green channel for the protein of interest and average intensity was measured. The red image was used to facilitate identification of cell boundaries and ROI’s were drawn freehand around clusters of cells. The cellular ROI’s were superimposed onto the green channel illuminating the protein of interest and average intensity was measured. The average intensity of the local background for each image was subtracted from the average intensity measurement of each ROI to determine the relative fluorescent units for each protein of interest.

Details regarding the number of experimental performed are included in all figure legends. All data are graphically represented as mean ± SD. Statistical analyses were performed using Prism software 10.0 GraphPad Software, Inc., San Diego, CA). A D’Agostino-Pearson normality test was performed to evaluate whether the data fit a Gaussian distribution. When the data were normally distributed, a two-tailed parametric T-test (n=2 groups) or a one-way ANOVA test (for ≥3 groups) was performed. When the data were not normally distributed, a Mann-Whitney test (n=2 groups) was performed. Of note, all data where ≥3 groups were compared were normally distributed. When present, the vehicle treatment condition served as the reference group for multiple comparisons analyses in the one-way ANOVA test. *p≦0.05. **p≦0.01. ***p≦0.001.

## RESULTS

### FTC, TFV, and DTG Differentially Cross the BBB

ART access to the CNS is an important public health concern as it contributes to maintenance of the brain as a viral reservoir and increases risk for neurologic sequelae, including cognitive and mood disorders in people living with HIV. While it is clear that ART enters the CNS compartment, albeit to a lower extent as compared to plasma and peripheral organs ^7, 97–108^, the precise mechanisms at the BBB that facilitate this remain poorly understood. To address this, we used primary human brain microvascular endothelial cells and primary human astrocytes to develop an *in vitro* model of the human BBB. In this system, endothelial cells and astrocytes express proteins present *in vivo*, notably the transferrin receptor, claudin-5, glucose transporter 1, VE-cadherin, occludin, PECAM-1, Zo-1, and GFAP (**Supplemental Figure 1**). Importantly, we demonstrated previously that this BBB model is dynamically regulated in response to inflammatory and angiogenic stimuli in an expected fashion ^91^.

The BBB model is generated by seeding endothelial cells into the upper, apical compartment while astrocytes grow on the basolateral underside for a period of three days until confluence is reached. During this time, the astrocytes extend their endfeet processes to make physical contact with the endothelia, effectively sealing the barrier ^87, 88^ (**Supplemental Figure 2A, 2C-F**). This model has high transendothelial electrical resistance and is impermeable to endogenous molecules excluded from an intact BBB *in vivo*, including inulin ^87, 88^ and albumin (**Supplemental Figure 2B**).

We used this BBB model previously to evaluate immune cell migration in the context of HIV ^86, 89, 90, 109^. Now, we leverage this system to evaluate the ability of three first-line ART drugs to cross the BBB: FTC, TFV, and DTG. Each ART drug (10 μM) was added to the apical portion of the model and allowed to pass to the basolateral chamber for 24 hours, after which time the media was collected and the concentration that passed determined by liquid chromatography/mass spectrometry. FTC and TFV readily crossed the BBB (**Figure 1**). However, FTC concentrations were considerably lower than TFV, at 792.2±136.3 versus 1183±142.9 ng/mL, respectively (p=0.0134, one-way ANOVA). In contrast, DTG was below the detectable limit of 100 ng/mL and therefore the concentration of drug that passed into the basolateral chamber was too low to quantitate (**Figure 1**). These findings demonstrate that, while ART can cross the BBB, differing drugs have distinct propensities to enter the CNS.

**Figure 1.**
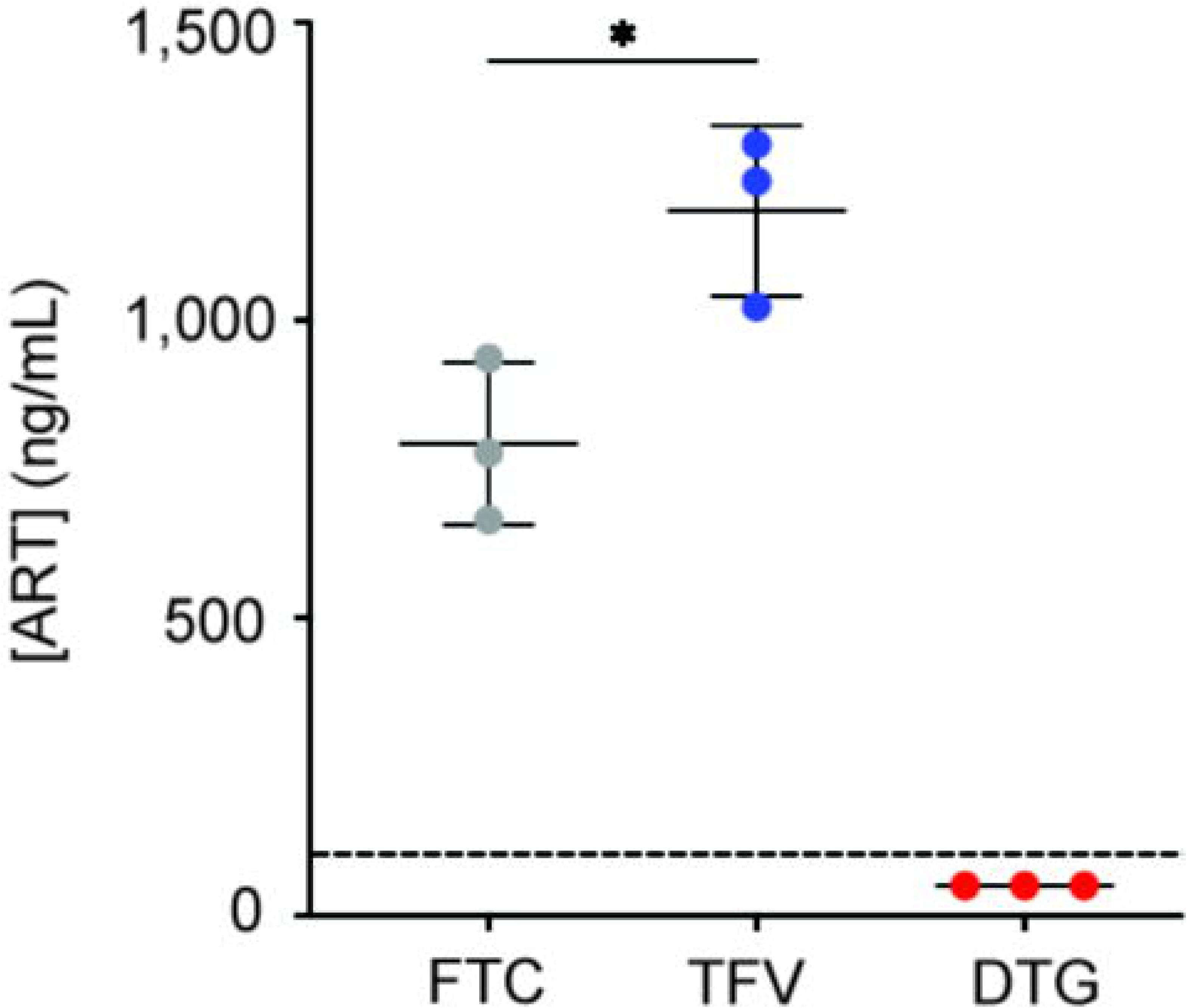
Differential ART Propensity to Cross the BBB. FTC (grey), TFV (blue), and DTG (red) (all at 10 μM) were added to the apical portion of the BBB model and allowed to extravasate into the basolateral portion for 24 hours at 37°C, 5% CO_2_. After this period of time, the media in the basolateral compartment was collected and the concentration of each ART drug that passed was measured by liquid chromatography/mass spectrometry. Dashed line denotes quantitative limit of detection for DTG. Three independent experiments (represented by individual dots), that included four technical replicates, were performed. Data are represented as mean ± standard deviation. *p<0.05. One-way ANOVA was performed.

### Cocaine Selectively Modulates FTC and TFV Extravasation Across the BBB

To evaluate the impact of comorbid substance use on ART availability in the CNS, BBB migrations were performed in the presence and absence of cocaine. The ability of ART to cross the BBB in the presence of LPS, a potent immune stimulus and inflammatory agent, was also performed as it is an important modulator of barrier function ^110–118^. FTC, TFV, and DTG (10 μM) were added to the apical portion of the BBB model in the presence and absence of cocaine (10 μM) or LPS (10 ng/mL) and allowed to pass to the basolateral chamber for 24 hours, after which time the media was collected and the ART concentration that passed determined by liquid chromatography/mass spectrometry. Cocaine increased the mean FTC concentration in the basolateral compartment by 268.4±6.5 ng/mL (p=0.0006, T-test, **Figure 2A**). While cocaine also impacted the concentration of TFV that passed the BBB, it had an opposing effect and decreased its presence by 293.2±26.7 ng/mL (p=0.0027, T-test, **Figure 2B**). Interestingly, LPS had an inconsistent effect (**Figure 2C-D**), where it caused a mean increase of 317.6±566.4 ng/mL for FTC (p=0.4339, T-test) and a decrease of 190.2±202.2 ng/mL for TFV (p=0.2449, T-test) that crossed the BBB. This suggests a specificity for cocaine’s impact on ART extravasation across the BBB, rather than a general mechanism that broadly occurs. The concentrations of DTG that passed the BBB in the presence of cocaine and LPS were below quantifiable limits (data not shown).

**Figure 2.**
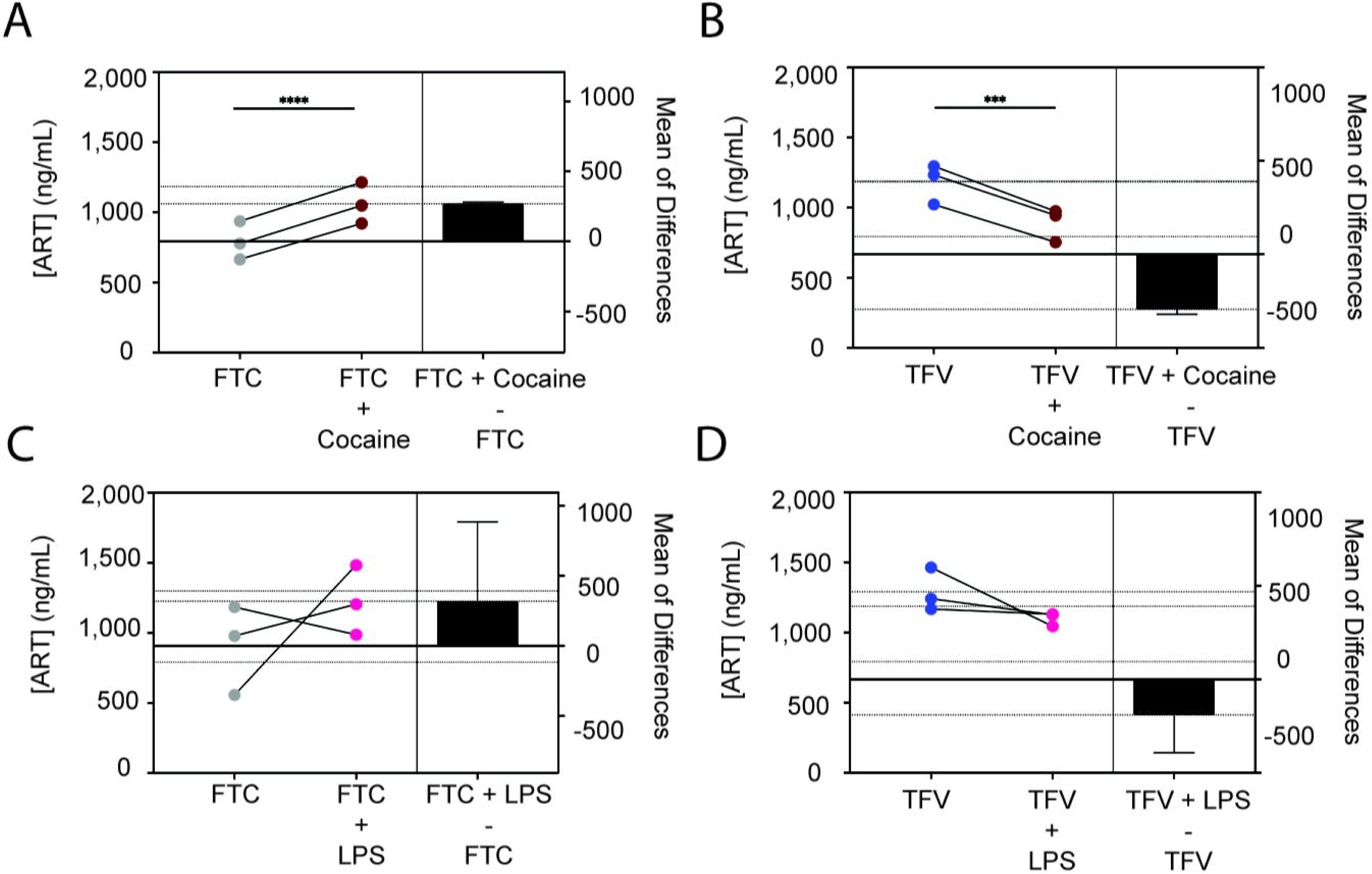
Cocaine, But Not LPS, Modulates ART Extravasation Across the BBB. FTC (grey, A and C) and TFV (blue, B and D) (both at 10 μM) were added to the apical portion of the BBB model and allowed to extravasate into the basolateral portion in the presence or absence of cocaine (10 μM, burgundy, A-B) or LPS (10 ng/mL, fuchsia, C-D) for 24 hours at 37°C, 5% CO_2_. The media in the basolateral compartment was collected and the concentration of each ART drug that passed was measured by liquid chromatography/mass spectrometry. Three independent experiments (represented by individual dots), that included four technical replicates, were performed. Estimation plots are shown where the left y-axis denotes ART concentration (ng/mL) and the right y-axis reflects the effect size (black bar), which is the difference between means of each condition. Data are represented as mean ± standard deviation. ***p<0.001. ****p<0.0001. Paired T-test was performed.

### Cocaine Does Not Impact BBB Permeability to Albumin Or Key Structural Endothelial Proteins

The differential selectivity by which FTC, TFV, and DTG crossed the BBB and the specificity for cocaine’s modulation of these processes, suggests well-regulated mechanisms are elicited that impact ART CNS concentrations. Nonetheless, it is important to consider that diminished BBB integrity may also occur, which would have an additional impact on ART extravasation. To test this possibility, we first evaluated BBB permeability to albumin, the most abundant plasma protein, which is unable to cross an intact BBB under homeostatic conditions. However, it can readily bypass breaches in a compromised BBB where it enters the CNS and contributes to pathology. As such, albumin is used clinically as an index of BBB damage.

We evaluated permeability of the BBB model to EBA, where albumin-conjugated Evans blue dye that passed to the basolateral chamber was quantitated spectrophotometrically (**Supplemental Figure 2B**). The BBB was treated with cocaine (10 μM), LPS (10 ng/mL), or vehicle control for 24 hours, after which time permeability to EBA was evaluated. To our surprise, BBB models treated with cocaine had only a slight increase in permeability (**Figure 3A**), as evidenced by a 50% increase (p=0.1244, one-way ANOVA) in the optical density at 620 nm (OD_620_), as compared to vehicle treatment. There was an even smaller impact of LPS on BBB permeability as treatment caused only a 13% increase in the OD_620_ (p=0.8387, one-way ANOVA), as compared to vehicle. In contrast, a complete breach of the BBB would have permitted all the EBA dye into the basolateral chamber and resulted in a 580% increase in the OD_620_ (p <0.0001, one-way ANOVA, **Figure 3A**), as compared to vehicle treatment.

**Figure 3.**
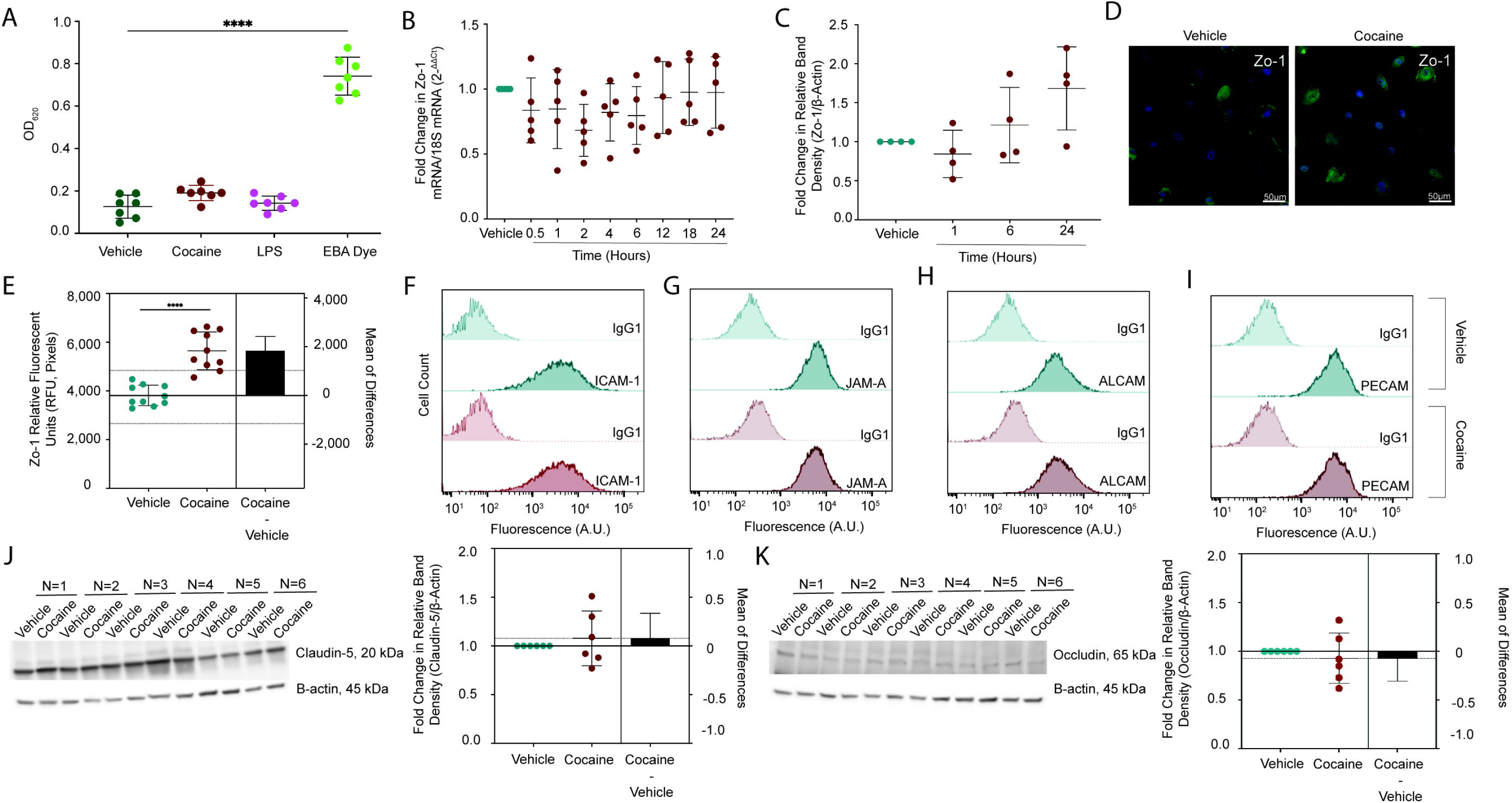
Cocaine Does Not Disrupt BBB Integrity. (A) BBB models were treated with cocaine (10 μM, burgundy), LPS (10 ng/mL, fuchsia), or vehicle (teal) for 24 hours, after which time permeability to EBA dye was performed. EBA was added to the apical portion of the BBB for 30 minutes at 37°C, 5% CO2, the media in the bottom collected, and evaluated spectrophotometrically at OD_620_. EBA dye alone (green) was used as a positive control to represent maximal BBB permeability. Seven individual experiments were performed (represented by individual dots). Data are represented as mean ± standard deviation. ****p<0.0001. One-way ANOVA was performed. (B) qRT-PCR was performed to evaluate Zo-1 mRNA following 0.5-24 hour treatment with cocaine (10 μM. burgundy). The 2-ΔΔCt method was performed to evaluate fold change in Zo-1 mRNA relative to 18S mRNA where vehicle treatment (teal) was set to 1. Five individual experiments were performed (represented by individual dots). Data are represented as mean ± standard deviation. (C) Western blot was performed to evaluate Zo-1 total protein expression following 1, 6, and 24 hour treatment with cocaine (10 μM, burgundy). Blots were stripped and reprobed to evaluate β-actin for protein normalization. The fold change in relative band intensity for Zo-1/β-actin was determined by densitometry where vehicle treatment (teal) was set to 1. Four individual experiments were performed (represented by individual dots). Data are represented as mean ± standard deviation. (D) Immunofluorescent microscopy was performed to evaluate Zo-1 (green) following treatment with cocaine (10 μM, right) or vehicle (left) for 24 hours. DAPI was used to visualize nucleus (blue). One paired representative image, out of 10 individual images, are shown. All scale bars = 50 μm. (E) Quantification of the fluorescent signal from Zo-1 immunofluorescent microscopy was performed for endothelial cells treated with cocaine (10 μM, burgundy) or vehicle (teal) for 24 hours. Ten independent experiments (represented by individual dots) were performed. Estimation plots are shown where the left y-axis denotes relative fluorescent intensity (RFU, pixels) and the right y-axis reflects the effect size (black bar), which is the difference between means of each condition. Data are represented as mean ± standard deviation. ****p<0.0001. Unpaired T-test was performed. (F-I) Flow cytometry was performed to evaluate cell surface expression of (F) ICAM-1, (G) JAM-A, (H), ALCAM, and (I) PECAM following 24-hour treatment with cocaine (10 μM, burgundy) or vehicle (teal). Fluorescence (arbitrary units) was evaluated for the specific protein of interest or following staining with an irrelevant, nonspecific isotype matched negative control antibody (IgG1). Data from one representative experiment, out of four individual experiments, are shown. (J-K) Western blot was performed to evaluate (J) claudin-5 and (K) occludin total protein expression in endothelial cells following 24-hour treatment with cocaine (10 μM, burgundy) or vehicle (teal). β-actin was used for protein normalization. Western blots demonstrating six independent experiments are shown (left). The fold change in relative band intensity for Zo-1/β-actin was determined by densitometry where vehicle treatment (teal) was set to 1 (right). Six independent experiments (represented by individual dots) were performed. Estimation plots are shown where the left y-axis denotes fold change in relative band intensity for the protein of interest relative to β-actin and the right y-axis reflects the effect size (black bar), which is the difference between means of each condition. Data are represented as mean ± standard deviation.

As albumin is a large tracer of 67 kDa that can only cross a substantially impaired BBB, we evaluated whether cocaine impacted the barrier integrity more subtly by diminishing key interendothelial junctions that promote transcellular integrity. First, we treated endothelial cells with cocaine (10 μM), or vehicle, between 0.5-24 hours and evaluated Zo-1 by qRT-PCR (**Figure 3B**). While there was, on average, a 20.4% decrease (p=0.2706-0.8724, one-way ANOVA) in Zo-1 mRNA between 0.5-6 hours, levels were restored to near basal levels by 24 hours (3% decrease in Zo-1 mRNA, p=0.9997, one-way ANOVA). A similar pattern occurred at the protein level evaluated by Western blot analysis (**Figure 3C**), where the relative band density for Zo-1 decreased at 1 hour (16% decrease, p=0.8944, one-way ANOVA), was later restored, and even trended towards an increase as compared to vehicle treatment by 24 hours (68% increase, p=0.0717, one-way ANOVA). As this was unexpected, we evaluated Zo-1 more comprehensively by immunofluorescent microscopy analysis following 24 hours of cocaine (10 μM) or vehicle treatment. Confirming the Western blot, cocaine induced a 48% increase (p <0.0001, T-test) in Zo-1 relative fluorescent unit (RFU) intensity after 24 hours of cocaine treatment, compared to vehicle (**Figure 3D-3E)**.

We next evaluated the impact of cocaine on additional tight junction and adhesion molecule proteins that serve to maintain BBB integrity by flow cytometry. Endothelial cells were treated with cocaine (10 μM) or vehicle for 24 hours, the cells gently removed from adherent culture with TrypLE to maintain surface antigens ^91^, and immunostained and analyzed by flow cytometry. Histogram plots, representative of four independent experiments, demonstrated that cocaine did not alter the cell surface expression of intercellular adhesion molecule 1 (ICAM-1), junctional adhesion molecule A (JAM-A), activated leukocyte cell adhesion molecule (ALCAM), and platelet-endothelial cell adhesion molecule (PECAM-1) (**Figure 3F-I**), as compared to vehicle treatment. In addition to the cell surface markers, we evaluated two tight junction molecules essential for BBB integrity, claudin-5 and occludin, by Western blot and determined that cocaine had an inconsistent and marginal impact on their expression with a 7% increase (p=0.5104, T-test) and 7% decrease (p=0.5128, T-test) in relative band density, respectively (**Figure 3J-K**).

Finally, we aimed to evaluate the global impact of cocaine on proteins involved in maintaining endothelial cell junctions. To accomplish this, we performed untargeted proteomics following 24 hours of treatment with cocaine (10 μM) or vehicle. Of the 4,831 identified proteins, cocaine modulated only 12 proteins relating to BBB integrity (**Table 2**). Of these, the most substantially impacted was carcinoembryonic antigen-related cell adhesion molecule 1 (CEACAM1), which cocaine induced a 78% increase in copy number (p=0.0035, T-test) and a 97% increase in intracellular concentration (p=0.0010, T-test), as compared to vehicle. The impact on CEACAM1 was atypical, however, as cocaine had a smaller impact on the remaining 11 proteins that ranged from a 6-40% change in copy number and an 8-55% change in intracellular concentration (**Table 2**). Together, these data suggest that in our system, cocaine does not substantially impact proteins involved in maintaining BBB integrity, does not increase permeability, and that its impact on ART extravasation into the CNS occurs through other mechanisms.

**Table 2.** Protein copy number and concentration following 24-hour treatment with cocaine (10 μM) or vehicle as determined by proteomics analysis. Mean ± standard deviation are shown. Asterisks indicate statistical significance in cocaine treated cells relative to vehicle. *p<0.05. **p<0.01. ***p<0.001. ****p<0.0001. Paired T-test. COL6A1, Collagen Type VI Alpha 1 Chain. CTSB, Cathepsin B. HPX, Hemopexin. MIF, Macrophage Migration Inhibitory Factor. PICALM, Phosphatidylinositol Binding Clathrin Assembly Protein. NRP2, Neuropilin 2. VTN, Vitronectin. VWF, Von Willebrand Factor.

|  | Vehicle |  | Cocaine |  |
| --- | --- | --- | --- | --- |
| Protein | Copy Number | Concentration (nM) | Copy Number | Concentration (nM) |
| CEACAM1 | 59,626 ±10,789 | 43.9 ± 7.5 | 106,220 ±<br>7,458** | 86.6 ± 4.4** |
| COL6A1 | 206,854 ±<br>12,118 | 152.2 ± 3.7 | 131,021 ±<br>14,120** | 106.9 ± 11.7** |
| CTSB | 8,484,178 ±<br>416,822 | 6,252.0 ± 355.4 | 6,593,383 ±<br>408,852** | 5,377.0 ± 230.0* |
| HPX | 6,543,012 ±<br>138,928 | 4,821.0 ± 139.0 | 4,814,035 ±<br>116364**** | 3,931.0 ± 184.8** |
| ICAM | 676,996 ±<br>14,380 | 499.0 ± 24.0 | 721,746 ±<br>23,606* | 588.8 ± 7.0** |
| MIF | 3,493,966 ±<br>119,612 | 2,573.0 ± 39.4 | 4,902,856 ±<br>535,590* | 3,996.0 ± 358.4** |
| PECAM | 1,202,038 ±<br>59,972 | 885.6 ± 46.24 | 956,743 ±<br>10,063** | 780.8 ± 9.6* |
| PICALM | 1,014,023 ±<br>21,188 | 747.1 ± 20.5 | 788,182 ±<br>30,189*** | 643.2 ± 25.7** |
| NRP2 | 287,851 ±<br>21,441 | 212.0 ± 13.9 | 186,651 ±<br>2,301** | 152.3 ± 3.6** |
| VTN | 1,208,330 ±<br>68,554 | 890.8 ± 65.2 | 862,826 ±<br>6,116*** | 704.3 ± 20.8** |
| VWF | 584,018 ±<br>20,536 | 430.1 ± 9.7 | 484,891 ±<br>17,129** | 395.6 ± 11.0* |

### Cocaine Inhibits PXR, the Master Regulator of Drug Transporters and Metabolizing Enzymes

As cocaine did not impact BBB integrity, we sought to evaluate the mechanisms by which it affected ART CNS access. We hypothesized that cocaine regulated cellular processes contributing to drug transport and metabolism at the BBB. To address this hypothesis, we turned our attention to PXR and CAR: transcription factors that serve key roles in regulating drug transport and metabolism following induction by xenobiotics in efforts to detox the cell (**Figure 4A**). We first performed Western blot to evaluate total protein levels in cell lysates obtained following 24-hour treatment with cocaine (10 μM) or vehicle and found that cocaine caused a 23% decrease in the relative band intensity for PXR (p<0.0001, T-test), as compared to vehicle (**Figure 4B, 4D**). Interestingly, this effect was specific to PXR as cocaine induced only a 7% decrease in the relative band intensity for CAR (p=0.3133, T-test, **Figure 4C, 4E**). We confirmed these findings by immunofluorescence and found a similar cocaine-induced decrease in PXR (24% decrease in fluorescent signal, p<0.0001, T-test, **Figure 4F, 4H**), which did not occur for CAR (1% decrease in fluorescent signal, p=0.9081, T-test, **Figure 4G-4I**). Of note, the cocaine-mediated decrease of PXR occurred in a dose-independent fashion (**Figure 5A**). All subsequent experiments were performed at a cocaine concentration of 10 μM.

**Figure 4.**
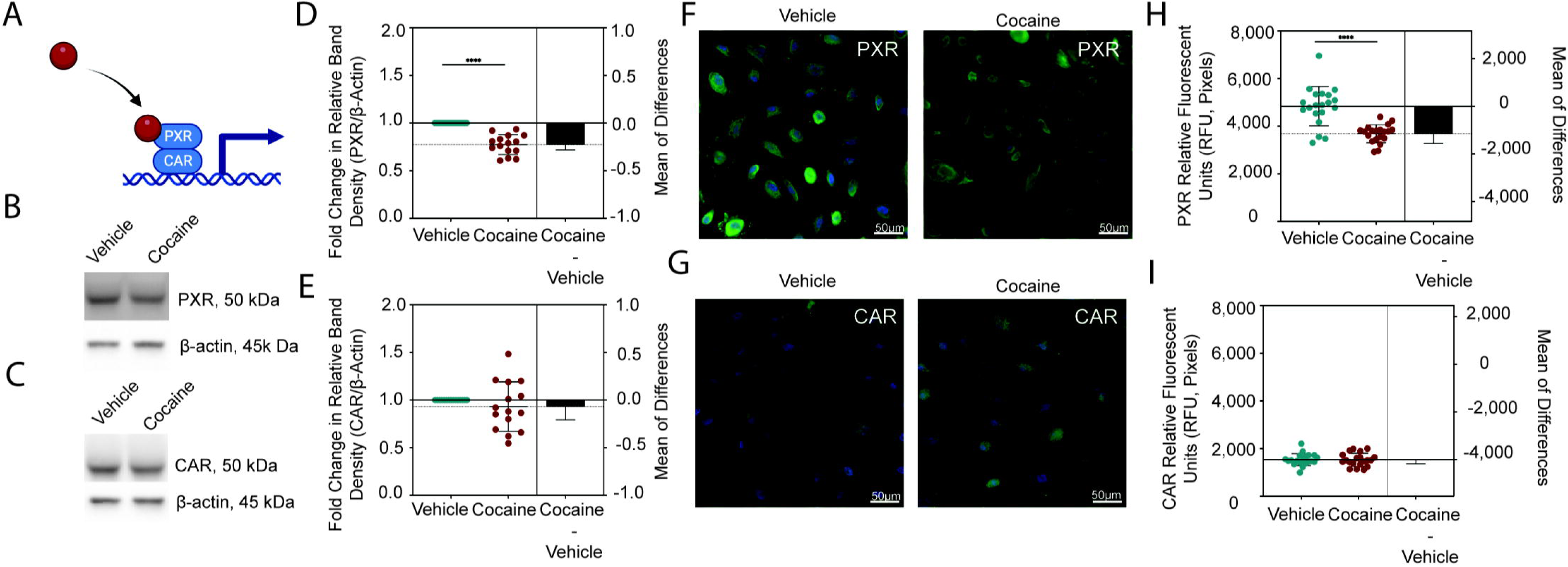
Cocaine Decreases PXR, But Not CAR, in Endothelial Cells. (A) Schematic representation depicting the transcriptional activity of PXR and CAR following ligand binding. (B-C) Western blot was performed to evaluate (B) PXR and (C) CAR following 24-hour treatment with cocaine (10 μM) or vehicle. β-actin was used for protein normalization. One western blot, representative of 15 independent experiments, is shown. (D-E) The fold change in relative band intensity for (D) PXR/β-actin and (E) CAR/β-actin was determined by densitometry where vehicle treatment (teal) was set to 1 (right). Fifteen independent experiments (represented by individual dots) were performed. Estimation plots are shown where the left y-axis denotes fold change in relative band intensity for the protein of interest relative to β-actin and the right y-axis reflects the effect size (black bar), which is the difference between means of each condition. Data are represented as mean ± standard deviation. ****p<0.0001. Unpaired T-test was performed. (F-G) Immunofluorescent microscopy was performed to evaluate (F) PXR or (G) CAR (green) following treatment with cocaine (10 μM, right) or vehicle (left) for 24 hours. DAPI was used to visualize nucleus (blue). One paired representative image, out of 20 individual images, are shown. All scale bars = 50 μm. (H-I) Quantification of the fluorescent signal from (H) PXR and (I) CAR immunofluorescent microscopy was performed for endothelial cells treated with cocaine (10 μM, burgundy) or vehicle (teal) for 24 hours. Twenty independent experiments (represented by individual dots) were performed. Estimation plots are shown where the left y-axis denotes relative fluorescent intensity (RFU, pixels) and the right y-axis reflects the effect size (black bar), which is the difference between means of each condition. Data are represented as mean ± standard deviation. ****p<0.0001. Unpaired T-test was performed.

**Figure 5.**
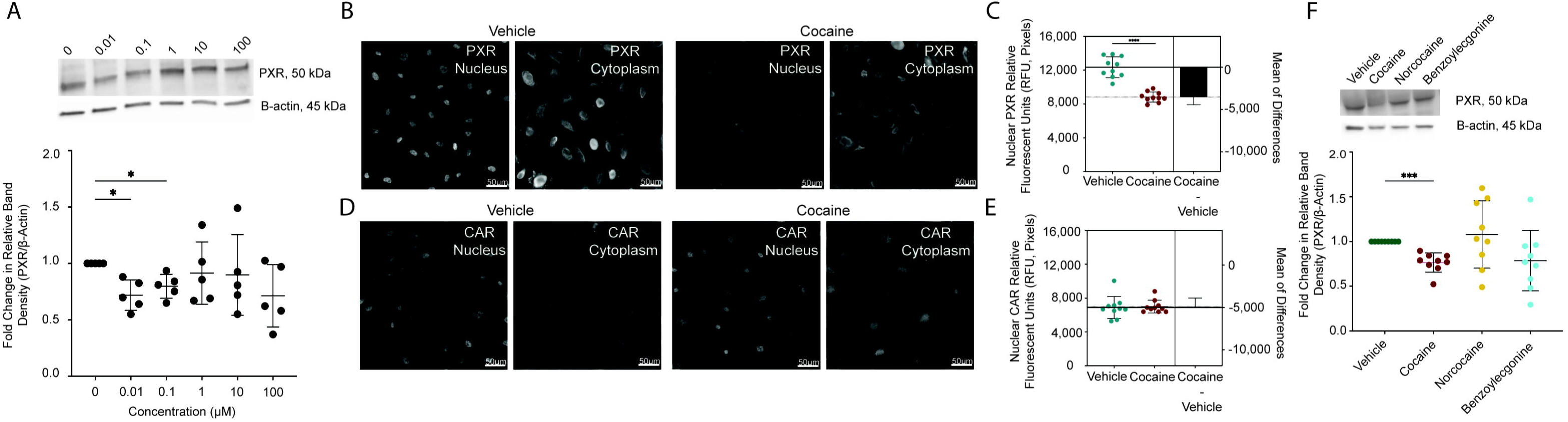
Cocaine’s Effect on PXR is Specific, Dose-Independent, and Occurs Primarily in the Nucleus. (A) Western blot was performed to evaluate PXR following 24-hour treatment with cocaine (0.01-100 μM) or vehicle (0 μM). β-actin was used for protein normalization. One western blot, representative of five independent experiments, is shown (top). The fold change in relative band intensity for PXR/β-actin was determined by densitometry where vehicle treatment was set to 1. Five independent experiments (represented by individual dots) were performed. The fold change in relative band intensity for PXR relative to β-actin is depicted (bottom). Data are represented as mean ± standard deviation. *p<0.05. One-way ANOVA was performed. (B-E) Immunofluorescent microscopy was performed to evaluate (B) PXR or (D) CAR following treatment with cocaine (10 μM, right) or vehicle (left) for 24 hours. The fluorescent signal for each respective protein that colocalized with DAPI was separated from that which occurred in the cytoplasm to facilitate analysis of PXR and CAR specifically in the nucleus. One paired representative image, out of 20 individual images, are shown. All scale bars = 50 μm. (C, E) Quantification of the nuclear fluorescent signal from (C) PXR and (E) CAR immunofluorescent microscopy was performed for endothelial cells treated with cocaine (10 μM, burgundy) or vehicle (teal) for 24 hours. Twenty independent experiments (represented by individual dots) were performed. Estimation plots are shown where the left y-axis denotes relative fluorescent intensity (RFU, pixels) of the nucleus and the right y-axis reflects the effect size (black bar), which is the difference between means of each condition. Data are represented as mean ± standard deviation. ****p<0.0001. Unpaired T-test was performed. (F) Western blot was performed to evaluate PXR following 24-hour treatment with cocaine (10 μM), its minor metabolite norcocaine (10 μM), its major metabolite benzoylecgonine (10 μM) or vehicle. β-actin was used for protein normalization. One western blot, representative of 9 independent experiments, is shown (top). The fold change in relative band intensity for PXR/β-actin was determined by densitometry where vehicle treatment was set to 1. Nine independent experiments (represented by individual dots) were performed. The fold change in relative band intensity for PXR relative to β-actin is depicted following cocaine (burgundy), norcocaine (yellow), benzoylecgonine (turquoise), or vehicle (teal) treatment (bottom). Data are represented as mean ± standard deviation. ***p<0.001. One-way ANOVA was performed.

We next evaluated the nuclear presence of PXR and CAR by immunofluorescent microscopy, as their transcriptional regulatory functions require translocation to the nucleus to affect drug transport and metabolism genes. The fluorescent signal for each respective protein that colocalized with DAPI was separated from that which occurred in the cytoplasm to facilitate analysis of PXR and CAR specifically in the nucleus. Interestingly, the PXR fluorescent signal in vehicle treated cells was higher in the nucleus (**Figure 5B-E**), as compared to that which occurred in the entire cell (**Figure 4F-I**), having a mean RFU of 12,349±1,222 versus 4,840±822, respectively. A similar effect occurred for CAR where the nuclear RFU was 6,896±1,302 while it was only 1,539±249.7 in the cytoplasm. Similar to that which occurred in the entire cell (**Figure 4H**), cocaine decreased the nuclear PXR RFU by 29% (p<0.0001, T-test) while having a 1% (p=0.7959, T-test) increase in nuclear CAR RFU (**Figure 5C, 5E**).

We next wanted to evaluate the specificity of the cocaine-mediated decrease in PXR. To accomplish this, we treated endothelial cells with cocaine, its minor metabolite norcocaine (10 μM), its major metabolite benzoylecgonine (10 μM), or vehicle for 24 hours and evaluated PXR by Western blot. As before, cocaine caused a 24% decrease (p=0.0005, one-way ANOVA) in the relative band intensity of PXR (**Figure 5F**). However, this did not occur for its metabolites. Indeed, norcocaine caused only an 8% increase (p=0.8612, one-way ANOVA) while benzoylecgonine had a 20% decrease (p= 0.2093, one-way ANOVA) in the relative band intensity of PXR. While benzoylecgonine’s effects on PXR were most similar to cocaine, they occurred inconsistently and had a large standard deviation of 34%.

### Cocaine Regulates Drug Transporter Expression and Activity

Cocaine’s modulation of PXR, but not CAR, has implications for drug transport across the BBB. To characterize this further, we evaluated the impact of cocaine on ten drug transporters known to interact with ART, or whose substrate structural similarity indicates the potential to impact ART tissue availability. We focused on five influx transporters, as well as five transporters involved in efflux, and determined that cocaine modulated eight of the ten proteins (**Figure 6**). Overall, cocaine decreased drug transporter expression, as compared to vehicle, where it promoted a loss of RFU for BCRP (18%, p=0.0166, T-test), ENT1 (49%, p<0.0001, T-test), MRP4 (23%, p=0.0006, T-test), OAT1 (45%, p<0.0001, T-test), OAT3 (17%, p<0.0001, T-test), OATP1A2 (24%, p<0.0001, T-test), and P-gp (24%, p<0.0001, T-test). OATP2A1 was the only evaluated transporter that had an increased RFU (66%, p<0.0001, T-test) following cocaine treatment.

**Figure 6.**
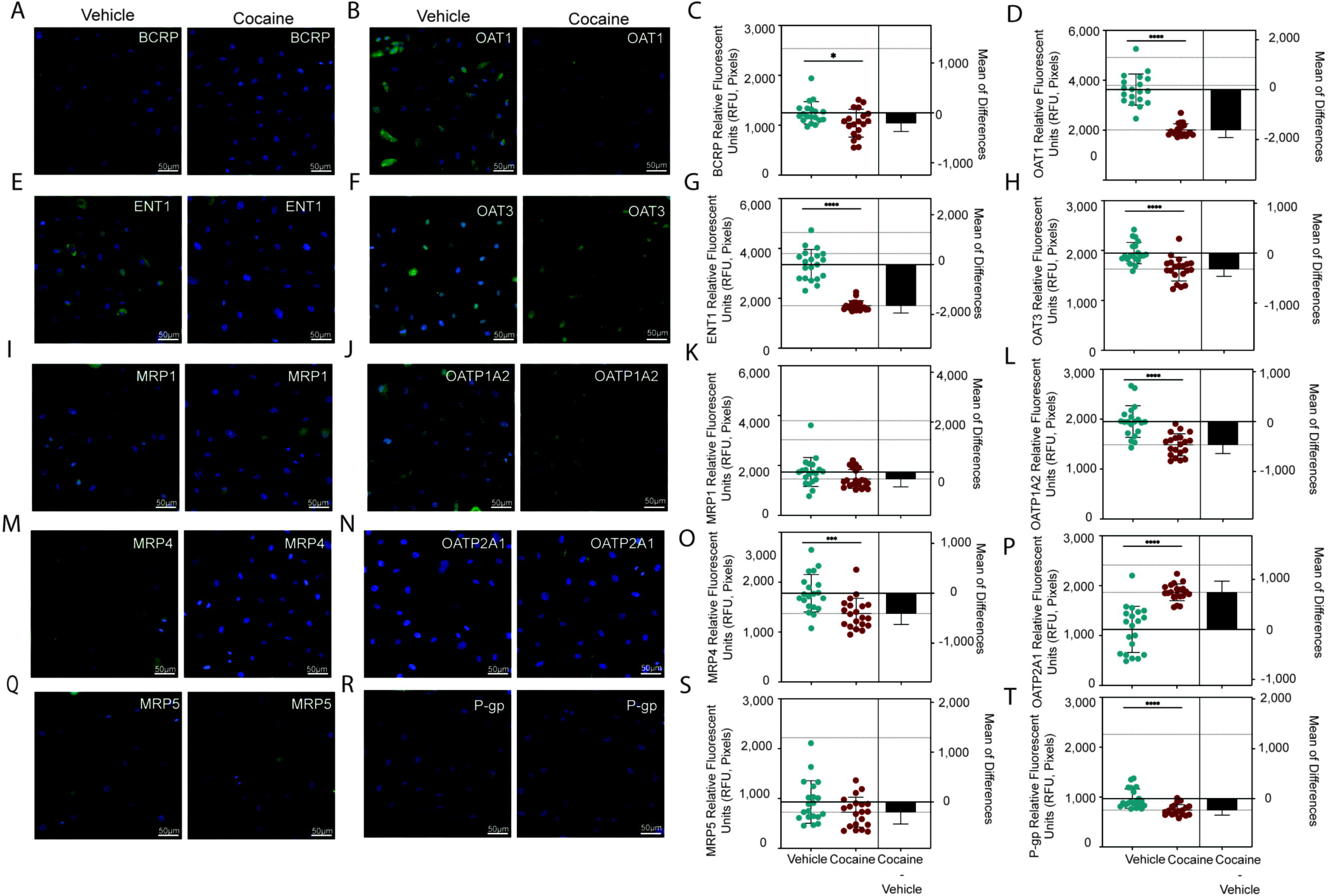
Cocaine Modulates Drug Transporter Expression. Immunofluorescent microscopy was performed to evaluate (A) BCRP, (B) OAT1, (E) ENT1, (F) OAT3, (I) MRP1, (J) OATP1A2, (M) MRP4, (N) OATP2A1, (Q) MRP5, or (R) P-gp (green) following treatment with cocaine (10 μM, right) or vehicle (left) for 24 hours. DAPI was used to visualize nucleus (blue). One paired representative image, out of 20 individual images, are shown. All scale bars = 50 μm. (C, D, G, H, K, L, O, P, S, T) Quantification of the fluorescent signal from immunofluorescent microscopy was performed for endothelial cells treated with cocaine (10 μM, burgundy) or vehicle (teal) for 24 hours. Twenty independent experiments (represented by individual dots) were performed. Estimation plots are shown where the left y-axis denotes relative fluorescent intensity (RFU, pixels) and the right y-axis reflects the effect size (black bar), which is the difference between means of each condition. Data are represented as mean ± standard deviation. *p<0.05. ***p<0.001. ****p<0.0001. Unpaired T-test was performed.

Intrigued by the implications of cocaine modulating ART transport across the BBB, we next sought to evaluate whether there were functional consequences for the altered presence of the influx and efflux proteins. We focused on three efflux transporters known to interact with ART that are modulated by PXR: BCRP, MRP4, and P-gp. To accomplish this, we pre-treated endothelial cells for 24 hours with cocaine, a specific inhibitor for each transporter, or the specific PXR inhibitor resveratrol.

Specifically, we used the BCRP inhibitor fumitremorgin (10 μM), the MRP4 inhibitor ceefourin 1 (10 μM), and P-gp inhibitor ritonavir (10 μM). Following treatment, the cells were loaded with a fluorescent dye (Hoechst 33342 for BCRP, monobromobimane for MRP4, and rhodamine 123 for P-gp) whose efflux is known to be mediated by our proteins of interest and compared the remaining intracellular fluorescent signal in the presence and cocaine and the inhibitors. We determined that cocaine inhibited the efflux activity of all three transporters, indicated by increased intracellular fluorescence (**Figure 7**). Each fluorescent dye rapidly entered the cell and was effluxed out after four hours, denoted by the loss of fluorescence when comparing Hoechst 33342 (**Figure 7A**), monobromobimane (**Figure 7B**), and rhodamine 123 (**Figure 7C**) to the vehicle condition. However, cocaine restored the fluorescent signal of all three dyes, indicating an inhibition of efflux out of the cell. Cocaine had a 39% increase (p=0.0162, T-test), a 91% increase (p=0.0011, T-test), and a 34% increase (p=0.0148, T-test) in the mean fluorescence intensity (MFI) attributed to Hoechst 33342, monobromobimane, and rhodamine 123, respectively. Interestingly, this diminished efflux activity mirrored the decreased expression of BCRP, MRP4, and P-gp induced by cocaine (**Figure 6C, 6O, 6T**). The impact of cocaine on the efflux transporters was comparable to that of their known inhibitors, strengthening the implications of cocaine in modulating the functional capacity of each transporter. Furthermore, resveratrol had a comparable inhibition on efflux activity as cocaine, demonstrating the importance of PXR in modulating transporter activity. These findings indicate that cocaine modulates transporter activity and identifies PXR as an important mechanism by which it alters the CNS efficacy of ART.

**Figure 7.**
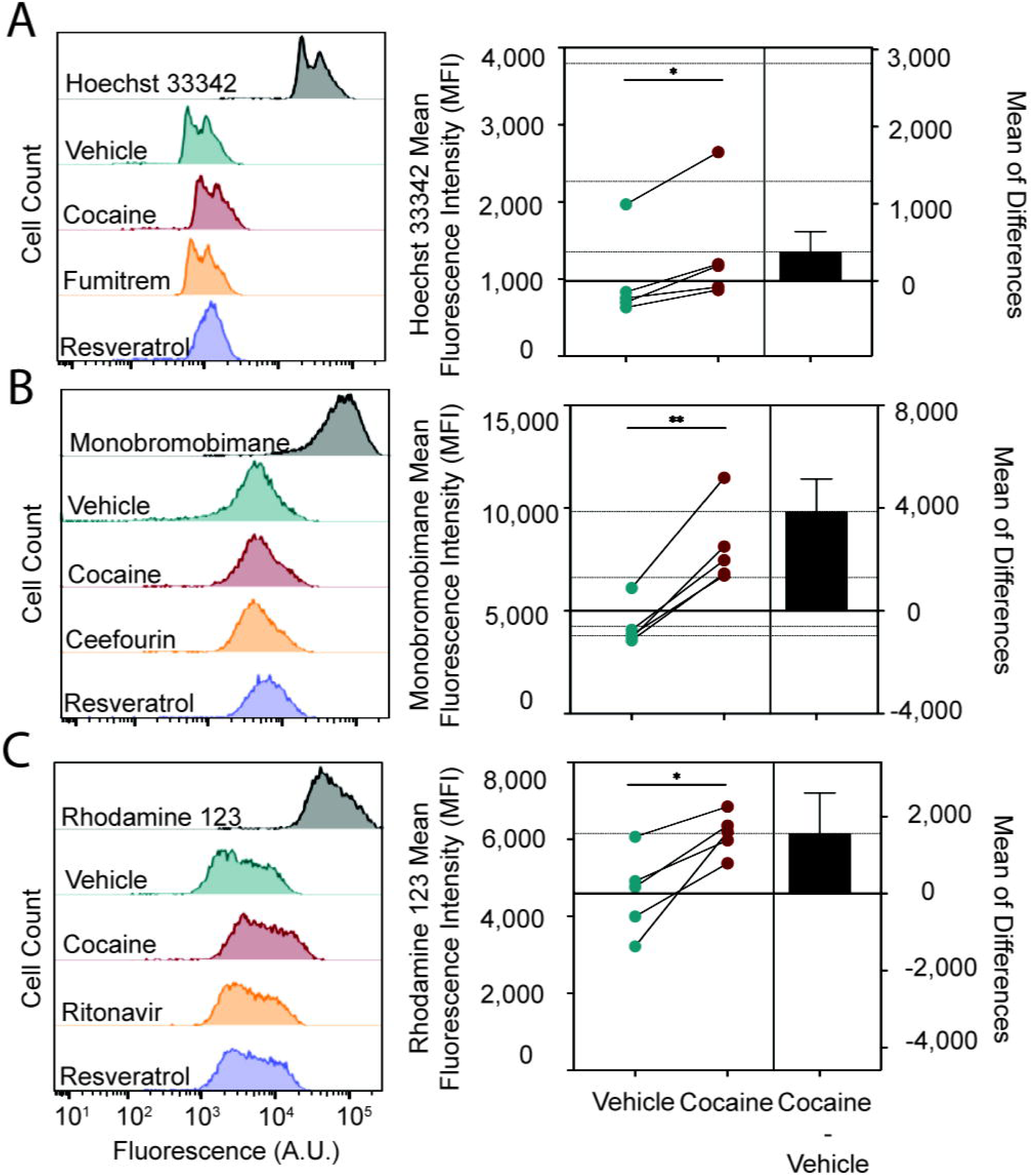
Cocaine Increases BCRP, MRP4, and P-gp Transport Activity. Endothelial cells were loaded with dyes specific for (A) BCRP (Hoechst 33342, 5 µg/mL), (B) MRP4 (monobromobimane, 10 μM), and (C) P-gp (rhodamine 123, 10 μM) for 1 hour (grey) and the dyes allowed to efflux out of the cell for four hours following pre-treatment with cocaine (10 μM, burgundy) or vehicle (teal). The cells were also pre-treated with specific inhibitors of (A) BCRP (10 μM, fumitremorgin), (B) MRP4 (10 μM, ceefourin 1), (C) P-gp (10 μM, ritonavir) (yellow) or PXR (resveratrol, 10 μM, lavender). Flow cytometric analysis was performed to evaluate the fluorescence and one histogram, representative of five independent experiments is shown (left). The fluorescent signal from flow cytometry was determined for endothelial cells pre-treated with cocaine (10 μM, burgundy) or vehicle (teal). Five independent experiments (represented by individual dots) were performed. Estimation plots are shown where the left y-axis denotes the mean fluorescent intensity (MFI, pixels) for (A) Hoechst 33342 (B) monobromobimane, and (C) rhodamine 123 and the right y-axis reflects the effect size (black bar), which is the difference between means of each condition (right). Data are represented as mean ± standard deviation. *p<0.05. **p<0.01. Paired T-test was performed.

### Cocaine Regulates Enzymes Involved in ART Metabolism and Biotransformation

In addition to its role in drug transport, PXR also contributes to drug metabolism by regulating phase I oxidative enzymes and phase II enzymes involved in glucuronic acid conjugation. CYP3A4 is one of the phase I enzymes of relevance for HIV treatment that is present at the BBB and may influence CNS ART availability. Of relevance for this study, CYP3A4 facilitates DTG metabolism into metabolite 3 ^119^, which was of interest as we were unable to quantify DTG’s ability to cross the BBB. Thus, we evaluated the impact of cocaine on CYP3A4 expression in endothelial cells following 24-hour treatment through Western Blot and immunofluorescent microscopy. Cocaine decreased the total protein levels of CYP3A4 as there was a 21% decrease (p <0.0001, T-test) in its relative band intensity, as compared to vehicle. This was confirmed microscopically where cocaine decreased the CYP3A4 RFU, relative to vehicle, by 23% (p=0.0035, T-test). Next, we evaluated the functional consequences of decreased CYP3A4 by evaluating its metabolic activity. Endothelial cells were pre-treated with cocaine or vehicle for 24 hours, loaded with the fluorogenic CYP3A4 substrate, BFC ^120–122^, and fluorescent signal measured over 80 minutes to evaluate formation of the fluorescent product, HFC. Cells were also pre-treated with rifampicin, as a positive control, and the PXR inhibitor resveratrol.

The rate of HFC production, indicated by fluorescent signal, occurred rapidly in the first 20 minutes after which time it began to plateau and eventually decline (**Supplemental Figure 3**). We used the linear portion of the curve in this first 20 minutes (**Figure 8E**) to evaluate the rate at which CYP3A4 converted BFC to HFC, termed CYP3A4 velocity. Cocaine increased the CYP3A4 velocity by 273% (p=0.0112, one-way ANOVA), as compared to vehicle, from 545±372 RFU/minute to 1,490±965 RFU/minute. This effect of cocaine was comparable to that of rifampicin, a well-known and clinically relevant CYP3A4 inducer, which increased the CYP3A4 velocity by 374% (p=0.0001, one-way ANOVA) to 2,039±1,125 RFU/minute. Of importance, resveratrol had the most profound effect on CYP3A4 by increasing its velocity by 585% to 3,188±1,613 RFU/minute, confirming the importance of PXR in CYP3A4 regulation.

**Figure 8.**
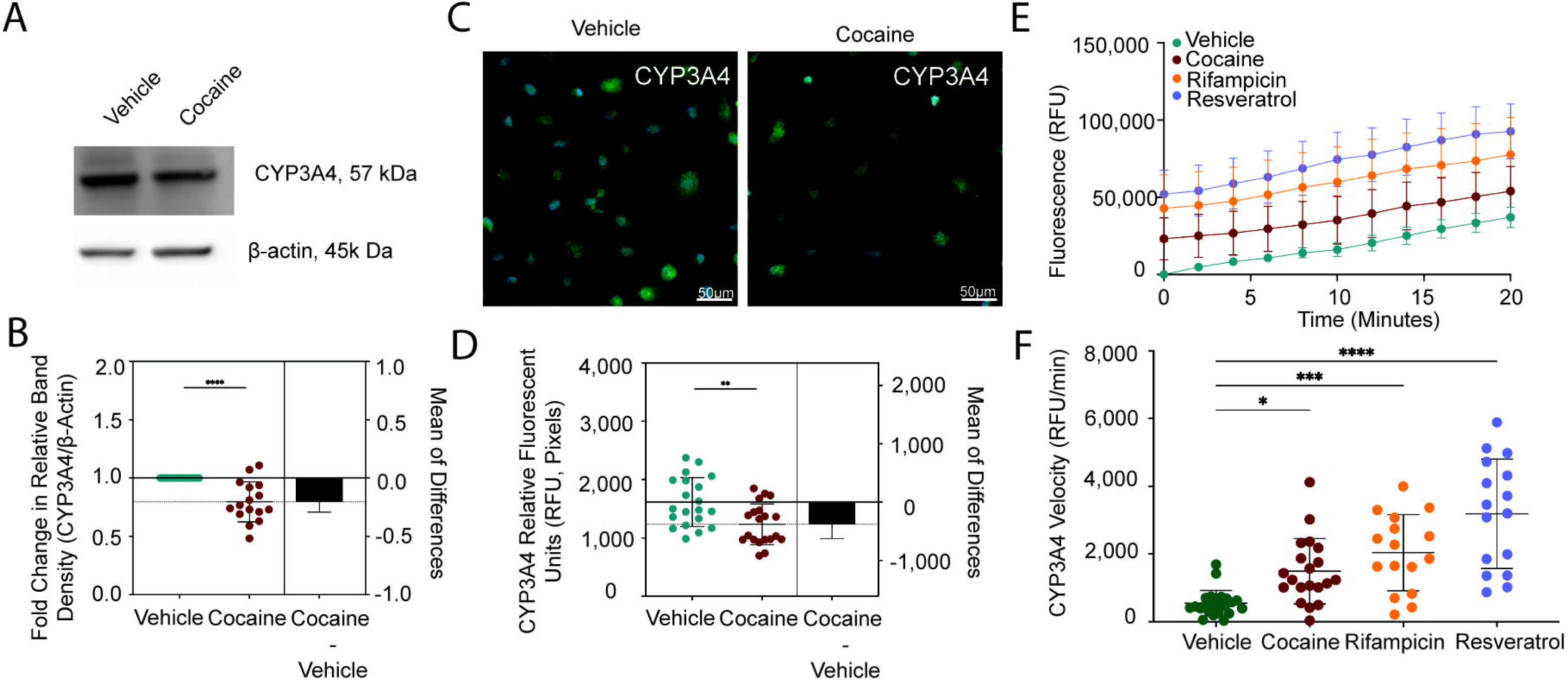
Cocaine Decreases CYP3A4 to Compensate for Increased Enzymatic Activity. (A) Western blot was performed to evaluate CYP3A4 following 24-hour treatment with cocaine (10 μM) or vehicle. β-actin was used for protein normalization. One western blot, representative of 16 independent experiments, is shown. (B) The fold change in relative band intensity for CYP3A4/β-actin was determined by densitometry where vehicle treatment was set to 1. Sixteen independent experiments (represented by individual dots) were performed. The fold change in relative band intensity for CYP3A4 relative to β-actin is depicted (bottom). Data are represented as mean ± standard deviation. *p<0.05. One-way ANOVA was performed. (C) Immunofluorescent microscopy was performed to evaluate CYP3A4 (green) following treatment with cocaine (10 μM, right) or vehicle (left) for 24 hours. DAPI was used to visualize nucleus (blue). One paired representative image, out of 20 individual images, are shown. All scale bars = 50 μm. (D) Quantification of the fluorescent signal from CYP3A4 immunofluorescent microscopy was performed for endothelial cells treated with cocaine (10 μM, burgundy) or vehicle (teal) for 24 hours. Twenty independent experiments (represented by individual dots) were performed. Estimation plots are shown where the left y-axis denotes relative fluorescent intensity (RFU, pixels) and the right y-axis reflects the effect size (black bar), which is the difference between means of each condition. Data are represented as mean ± standard deviation. **p<0.01. Unpaired T-test was performed. (E) Endothelial cells were pre-treated with cocaine (10 μM, burgundy), rifampicin (1 μM, yellow), resveratrol (10 μM, lavender), or vehicle (teal) for 24 hours, after which time the cells were loaded with BFC (2 μM). The enzymatic capacity of CYP3A4 to convert BFC to HFC was determined for the first 20 minutes as determined by fluorometric quantitation at excitation and emission wavelengths of 405/535 nm. Twelve independent experiments that contained eight technical replicates per condition were performed. Data are represented as mean ± standard deviation. **p<0.01. ****p<0.0001. Unpaired T-test was performed. (F) The rate at which BFC was converted to HFC is depicted as CYP3A4 velocity (RFU/min) for the earliest time points (2 and 4 minutes) to evaluate maximal enzymatic activity. The CYP3A4 velocity for each time point was pooled for both time points. Twelve independent experiments for each time point (represented by combined 24 individual dots) were performed. Data are represented as mean ± standard deviation. *p<0.05. ***p<0.001. ****p<0.0001. One-way ANOVA was performed.

We also wanted to evaluate the effect of cocaine on the metabolism of TFV and FTC, which are not CYP3A4 substrates. TFV and FTC are given as prodrugs that require biotransformation through a series of phosphorylation events to become pharmacologically capable of inhibiting HIV reverse transcriptase. Endogenous kinases, including adenylate kinases, are used to accomplish these phosphorylation events. Of the nine adenylate kinase isoforms, we evaluated AK1, AK2, and AK6 as they are the only isoforms that can use all ribonucleoside triphosphates as phosphate donors, as well as AK5 because it is exclusively expressed in the brain. Endothelial cells were treated with cocaine or vehicle for 24 hours and adenylate kinase expression was evaluated by immunofluorescent microscopy. Cocaine modulated AK1, AK5, and AK6, but not AK2, by causing a 20% increase (p=0.0012, T-test), a 13% decrease (p=0.0334, T-test), and a 19% increase (p=0.0011, T-test), in the RFU as compared to vehicle, respectively. Collectively, these findings demonstrate the implications of cocaine and PXR in altering ART metabolism that can impact its availability in the brain and ability to be efficacious in suppressing HIV.

## DISCUSSION

In this study, we demonstrated that cocaine modulates ART availability in the brain through regulation of drug transport and metabolism pathways at the BBB. Unexpectedly, we determined that cocaine did not perturb BBB integrity, but rather, downregulated PXR – the master regulator of drug transport and metabolism – to mediate its effects. Of importance, the impact of cocaine on ART extravasation across the BBB was not uniform, but instead, varied by drug. Our findings have profound implications for treatment of people with HIV with comorbid cocaine use disorders. Further, this work raises additional concerns for all substance using populations, not only those living with HIV, as the PXR-mediated drug transport and metabolism pathways explored in our present work are implicated in facilitating CNS access for therapeutics that treat essentially every neurologic disease.

The brain represents a major HIV reservoir. As such, the CNS efficacy of ART is a substantial barrier to HIV eradication efforts and remains a focus of much scientific and clinical investigation. Efforts to understand ART availability in the brain were once limited to predictive analyses based on chemical properties of individual drugs and their cerebrospinal fluid concentrations ^108, 123–125^. However, more recent evidence obtained from sampling distinct brain regions demonstrates that unexpectedly higher ART levels reach the CNS than previously considered ^7, 97–108, 126–128^. Even still, these concentrations are not comparable to those present in peripheral organs and people with HIV continue to have neurologic complications despite having undetectable plasma viral loads. Together, these concerns highlight the importance of understanding the molecular mechanisms by which ART traverses the BBB. Currently, our knowledge is limited and restricted to extrapolation from studies in peripheral organs – namely liver and small intestine. From these important studies we understand the transporters and metabolic enzymes capable of interacting with ART, including CYP3A4, P-gp, BCRP, and MRP4, as well as the importance of key transcriptional regulators PXR and CAR. As these proteins are also present and functional at the BBB, we assume that these same processes are involved in facilitating ART extravasation into the brain. However, studies demonstrating this are lacking. Our study addresses this gap in knowledge by demonstrating the importance of PXR in regulating drug transporter and metabolizing enzyme activity at the BBB.

Substance use is an important contributor to CNS HIV disease as it modulates neuroinflammatory, oxidative stress, and energy metabolic pathways ^60, 62, 64, 65, 129–134^. Additionally, there is a strong premise that substance use adversely impacts the BBB ^61, 66, 67, 135–146^. Most studies focused primarily on BBB integrity, transendothelial electrical resistance, and permeability. However, to our knowledge, there have been no studies evaluating the impact of substance use on ART’s ability to traverse the BBB into the CNS compartment. For these reasons, we sought to evaluate the impact of substance use on this important, but understudied, aspect that perpetuates HIV CNS disease using cocaine as a model illicit substance. We determined that cocaine increased FTC’s ability to cross the BBB, while decreasing that of TFV. This was intriguing because cocaine reversed their inherent differential ability to penetrate the BBB. Cocaine’s opposing effect on the ability of FTC and TFV to cross the BBB is quite striking, and concerning, as they are remarkably similar: they belong to the same ART class and have comparable molecular weights, structures, biophysical properties, pharmacokinetic distribution properties, substrate specificity, and phosphorylation-dependency to become pharmacologically capable of inhibiting HIV. This suggests that *in silico* and other predictive modeling analyses would be ineffective at predicting the differential impact of cocaine on ART availability in brain, which could have devastating clinical consequences. Our findings are an initial attempt at evaluating the impact of substance use on ART entry to the CNS and demonstrates that additional studies are warranted that consider the remaining ART drugs, as well as other substances of abuse. Furthermore, our findings provide caution to people without HIV who are prescribed these medications for pre-exposure prophylaxis (PrEP), as FTC and either the disoproxil fumarate or alafenamide formulations of TFV are co-administered in Truvada^®^ and Descovy^®^, respectively, the FDA approved drugs for HIV prevention ^147–150^. Indeed, care must be taken as our study raises the concern that cocaine use, and potentially other substances of abuse as well, while taking PrEP may alter its bioavailability and potential to prevent HIV transmission.

We were surprised to determine that cocaine didn’t adversely impact BBB integrity or permeability as it contrasts with the general consensus in the field. We posit that experimental differences in study design contribute to our discrepant finding, including species of origin, BBB model, and the tracers used to evaluate permeability. While unexpected, our analyses included measures of albumin permeability, tight junction and adhesion molecule expression, and evaluation of the entire proteomic landscape. Indeed, we employed qRT-PCR, Western blot, flow cytometry, and mass spectrometry to evaluate multiple aspects that help maintain BBB integrity. Thus, while unexpected, our complimentary methodologies are rigorous and demonstrate that cocaine’s impact on ART extravasation occurred by additional mechanisms. Our work provides evidence of the importance of evaluating BBB function beyond integrity and permeability measures by including functional analyses, such as drug extravasation studies. Further, it urges investigators to evaluate critically seemingly discrepant results if their disease model of interest does not perturb the BBB, as other underappreciated mechanisms may be involved. It is our hope that our findings expand the scope of considerations for how illicit substances, and other disease settings, impact BBB function.

Cocaine’s functional capacity to impact BBB transport pathways is of major interest in HIV CNS treatment strategies. The brain is notoriously difficult to treat as ∼98% of all therapeutics fail to penetrate the BBB ^151–155^. As a result, understanding the mechanisms by which therapeutics and illicit substances alter influx and efflux transport mechanisms is critical for effective treatment plans. We found that cocaine modulated eight transporters, where primarily decreased expression occurred (7/8 transporters). Five of these transporters facilitate influx into the brain, while the remainder are involved in efflux. Not only were the expression levels of the proteins affected, but cocaine also decreased their functional capacity to transport well-characterized substrates. Importantly, inhibiting PXR with resveratrol promoted an effect comparable to cocaine, further implicating the transcription factor as an important transport regulatory mechanism. These findings suggest that cocaine inhibits the potential of ART, and possibly other therapeutics as well, from entering the brain due to decreased influx transporter expression and activity. However, our work also indicates that cocaine also stabilizes CNS concentrations by preventing efflux out of the brain. These competing mechanisms provide insight regarding why cocaine may have distinct effects on the CNS concentrations of differing ART drugs. To our knowledge, this is the first time cocaine is implicated in modulating these processes. However, it should be noted that cocaine’s effect on drug transport shouldn’t be all that surprising, as it is classically known to mediate its rewarding effects by inhibiting dopamine, norepinephrine, and serotonin transporters ^156–168^. Further, there is increasing evidence that cocaine may impact other transporters, including those involved in glymphatic CNS clearance mechanisms ^169^. Interestingly, other addictive substances, including nicotine, opioids, and ethanol, affect transporter activity suggesting an overarching mechanism by which substance use may contribute to drug:drug interactions that adversely impact treatment strategies ^170–176^. Our findings posit that further investigation regarding the mechanisms by which cocaine, and other illicit substances, modulate CNS transport mechanisms more broadly are warranted, as they are integral in maintaining the specialized microenvironment of the brain that also have implications for therapeutic efforts.

We identified drug metabolism and biotransformation as additional mechanisms by which cocaine regulates the ability of ART to cross the BBB. Initially, we focused on CYP3A4 to provide insight into cocaine’s impact on DTG CNS concentrations, as we were unable to quantify its BBB transport. However, our findings have much broader implications as CYP3A4 is involved in metabolism of many ART drugs, including reverse transcriptase inhibitors, protease inhibitors, entry inhibitors, and other integrase inhibitors. Our findings demonstrate that cocaine decreases CYP3A4 expression by a PXR-mediated mechanism. Interestingly, this decrease in expression contributes to increased CYP3A4 enzymatic activity, as measured by HFC production. These findings suggest a compensatory mechanism by which PXR decreases CYP3A4 at the protein level to accommodate for its increased enzymatic efficiency. This impact of cocaine on CYP3A4 activity is remarkable as it is comparable to that of the well-known and clinically relevant CYP3A4 inducer, rifampicin. These findings have substantial clinical implications as CYP3A4 is the most abundant human cytochrome p450 isoform. Additionally, CYP3A4 is involved in the metabolism of 60% of all prescribed therapies. Our work identifies cocaine as a major modulator of CYP3A4, comparable to rifampicin, that has the potential to alter pharmacokinetics and dosing strategies for ART and other therapeutic drugs – including those that act in peripheral organs. This is the first time, to our knowledge, cocaine has been identified to regulate CYP3A4 enzymatic activity. Interestingly, other illicit substances also modulate CYP3A4 and other cytochrome P450’s ^170–176^. Furthermore, substance use can impact plasma concentrations of ART and viral rebound, independent of adherence concerns ^177–179^. Together, our findings demonstrate that additional attention is warranted in the clinical care of substance using populations with HIV, where measures of viral suppression and plasma ART concentrations are evaluated more regularly to evaluate treatment efficacy.

In addition to CYP3A4, we determined that cocaine also regulated the adenylate kinases involved in biotransformation of ART prodrugs into their pharmacologically active counterparts capable of suppressing HIV. Specifically, cocaine increased AK1, AK5, and AK6, but not AK2, demonstrating selectivity in its effects. As AK1 is ubiquitously expressed, our findings suggest cocaine’s impact on ART biotransformation may occur throughout the body, including the immune cells that are the primary target for HIV. Our findings also demonstrate a mechanism by which cocaine may specifically regulate ART CNS concentrations through modulation of the brain-specific AK5 isoform. This suggests a unique role for cocaine in regulating the ability of ART to be efficacious specifically in the CNS reservoir, primarily microglia, macrophages, and potentially astrocytes as their infection is a point of much discussion in the field.

## CONCLUSIONS

Our findings identify cocaine as an important contributor to the CNS efficacy of ART by altering its ability to cross the BBB and regulating PXR-mediated drug transport and metabolism pathways. Contrary to convention, cocaine’s effects did not breach BBB integrity, as evidenced by targeted evaluation of key proteins involved in barrier integrity, global proteomic evaluation, and albumin permeability measures. Further, cocaine increased FTC’s ability to cross the BBB while decreasing that of TFV, providing additional evidence of regulated, nuanced, and selective mechanisms rather than general loss of permeability. For the first time, we introduce awareness of the clinical ramifications of comorbid substance use in HIV cure strategies, specifically for viral eradication strategies in the brain. Furthermore, our findings provide insight into cocaine’s impacts on therapeutic strategies beyond HIV treatment, as PXR’s regulation of P-gp, BCRP, MRP4, and CYP3A4 activity are involved in drug disposition for numerous disorders.

## LIST OFF ABBREVIATIONS

(TBS-T): 1X Tris-Buffered Saline containing 0.1% Tween-20
(BFC): 7-benzyloxy-4-trifluoromethylcoumarin
(HFC): 7-hydroxy-4-trifluoromethylcoumarin
(ABC): ATP-binding cassette
(ART): Antiretroviral therapy
(AIDS): Acquired immunodeficiency syndrome
(ALCAM): Activated leukocyte cell adhesion molecule
(BBB): Blood brain barrier
(BCRP): Breast Cancer Resistant Protein
(CTSB): Cathepsin B
(CNS): Central nervous system
(COL6A1): Collagen Type VI Alpha 1 Chain
(cART): Combined ART
(M199C): Complete M199 media
(DPBS): Dulbecco’s phosphate-buffered saline without calcium or magnesium
(DTG): Dolutegravir
(EDTA): Ethylenediaminetetraacetic acid
(FTC): Emtricitabine
(ENT1): Equilitative nucleoside transporter
(EBA): Evans Blue dye conjugated to albumin
(FBS): Fetal bovine serum
(HPX): Hemopexin
(HIV): Human immunodeficiency virus-1
(ICAM-1): Intercellular Adhesion Molecule 1
(JAM-A): Junctional adhesion molecule A
(LPS): Lipopolysaccharide
(MIF): Macrophage Migration Inhibitory Factor
(MFI): P Mean fluorescence intensity
(M199): Medium 199
(MRP4): Multidrug resistance-associated protein 1
(MRP2): Multidrug resistance-associated protein 2
(MRP4): Multidrug resistance-associated protein 4
(NRP2): Neuropilin 2
(OAT1): Organic anion transporter 1
(OAT3): Organic anion transporter 3
(OATP1A2): Organic anion-transporting polypeptide 1A2
(OD620): Optical density at 620 nm
(PECAM-1): Platelet-endothelial cell adhesion molecule
(P-gp): P-glycoprotein
(PBS): Phosphate buffered saline
(ICALM): Phosphatidylinositol Binding Clathrin Assembly Protein
(PrEP): Pre-exposure prophylaxis
(PXR): Pregnane-X receptor
(ROI): Region of interest
(RFU): Relative fluorescent intensity
(SLC): Solute carrier
(TFV): transporter Tenofovir
(VWF): Vitronectin
(VWF): Von Willebrand Factor
(Zo-1): Zonula occludens-1

## SUPPLEMENTARY INFORMATION

**Supplemental Figure 1. Brain Microvascular Endothelial Cells and Astrocytes Express Characteristic Markers**. Immunofluorescent microscopy was performed to evaluate expression of anticipated markers in (A-G) primary human brain microvascular endothelial cells and (H) primary human astrocytes. (Left panels) Antibodies with specificity to (A) CD71, (B) claudin-5, (C) GLUT-1, (D) VE-Cadherin, (E) occludin, (F) PECAM-1, (G) Zo-1, and (H) GFAP were coupled to Alexa Fluor 488 for analysis. (Middle panels) DAPI was used to visualize nucleus. (Right panels) Merge depicts the combined signal for proteins of interest (green) and DAPI (blue). Representative images, out of 20 independent images, are shown. All scale bars = 50 μm.

**Supplemental Figure 2. I*n vitro* Model of the Human BBB.** (A) Schematic representation of our transwell BBB model where primary human brain microvascular endothelial cells are seeded on the upper, apical compartment and primary human astrocytes are seeded on the underside of a polycarbonate membrane with 3 μm pores in the basolateral compartment. (B) Schematic representation of albumin permeability assay, where EBA dye is added to the apical portion and permitted to pass to the basolateral side for 30 minutes at 37°C, 5% CO_2_. The media in the basolateral side is collected and spectrophometrically read at OD_620_ to evaluate BBB permeability. (C-F) The polycarbonate membrane from the BBB model was collected, immunostained, and immunofluorescent microscopy performed. (C-D) Wheat germ agglutinin (WGA) depicts cell morphology in red and (E-F) demonstrates expression of the astrocyte and endothelial cell markers GFAP and VE-Cadherin, respectively in green. DAPI was used to visualize nucleus (blue). Representative images, out of 3-5 independent images, are shown. All scale bars = 50 μm.

**Supplemental Figure 3. Complete Duration of CYP3A4 Metabolic Activity Assay**. Endothelial cells were pre-treated with cocaine (10 μM, burgundy), rifampicin (1 μM, yellow), resveratrol (10 μM, lavender), or vehicle (teal) for 24 hours, after which time the cells were loaded with BFC (2 μM). The enzymatic capacity of CYP3A4 to convert BFC to HFC was determined for 80 minutes as determined by fluorometric quantitation at excitation and emission wavelengths of 405/535 nm. Twelve independent experiments that contained eight technical replicates per condition were performed. Data are represented as mean ± standard deviation.

**Supplemental Figure 4. Cocaine Modulates AK1, AK5, and AK6 Expression**. Immunofluorescent microscopy was performed to evaluate (A) AK1, (B) AK2, (C) AK5, and (G) AK6 (green) following treatment with cocaine (10 μM, right) or vehicle (left) for 24 hours. DAPI was used to visualize nucleus (blue). One paired representative image, out of 20 individual images, are shown. All scale bars = 50 μm. Quantification of the fluorescent signal from immunofluorescent microscopy was performed for endothelial cells treated with cocaine (10 μM, burgundy) or vehicle (teal) for 24 hours. Twenty independent experiments (represented by individual dots) were performed. Estimation plots are shown where the left y-axis denotes relative fluorescent intensity (RFU, pixels) and the right y-axis reflects the effect size (black bar), which is the difference between means of each condition. Data are represented as mean ± standard deviation. *p<0.05. **p<0.01. Unpaired T-test was performed.

## DECLARATIONS

### Ethics approval and consent to participate

Not applicable

### Consent for publication

Not applicable

### Availability of data and materials

All data generated or analyzed during this study are included in this published article and are available from the corresponding author on reasonable request. All vendor original LCMS data and processed data has been made publicly available through the ProteomeXchange and MASSIVE public repositories^180^. Data can be directly accessed through FTP at the following link: ftp://MSV000092529@massive.ucsd.edu.

### Authors’ contributions

LBF, SK, ASP, RCC, AM, HW, BRF, and DWW performed experiments. LBF, SK, ASP, RCC, HW, BCO, and DWW and analyzed data. DWW was responsible for conceptualization of the study design. DWW wrote the original draft of the manuscript with review and editing from all authors. All authors read and approved the final manuscript.

## Acknowledgements

We thank Mr. Mark Marzinke of the Clinical Pharmacology Analytical Lab for his assistance with antiretroviral therapy determination. We acknowledge and thank the NIDA Drug Supply Program for providing the cocaine hydrochloride used in this study. We thank the members of the Johns Hopkins School of Medicine Retrovirus Laboratory in the Department of Molecular and Comparative Pathobiology for their support. Images in figures were created using Biorender.

