## Supplemental Figures for "Cocaine Regulates Antiretroviral Therapy CNS Access Through Pregnane-X Receptor-Mediated Drug Transporter and Metabolizing Enzyme Modulation at the Blood Brain Barrier"

^1^Department of Molecular and Comparative Pathobiology, ^2^Department of Neuroscience, Johns Hopkins School of Medicine, Baltimore, Maryland 21205, ^3^Department of Pharmacology and Molecular Sciences, ^4^Department of Medicine, Division of Clinical Pharmacology, Johns Hopkins School of Medicine, Baltimore, Maryland 21205, ^5^Department of Molecular Microbiology & Immunology, Johns Hopkins School of Public Health, Baltimore, Maryland 21205

*Address correspondence to**: Dionna W. Williams, Ph.D.**, Department of Molecular and Comparative Pathobiology, Johns Hopkins University School of Medicine, 733 N. Broadway St., Miller Research Building 833, Baltimore, MD 21205,

**SUPPLEMENTAL MATERIALS**

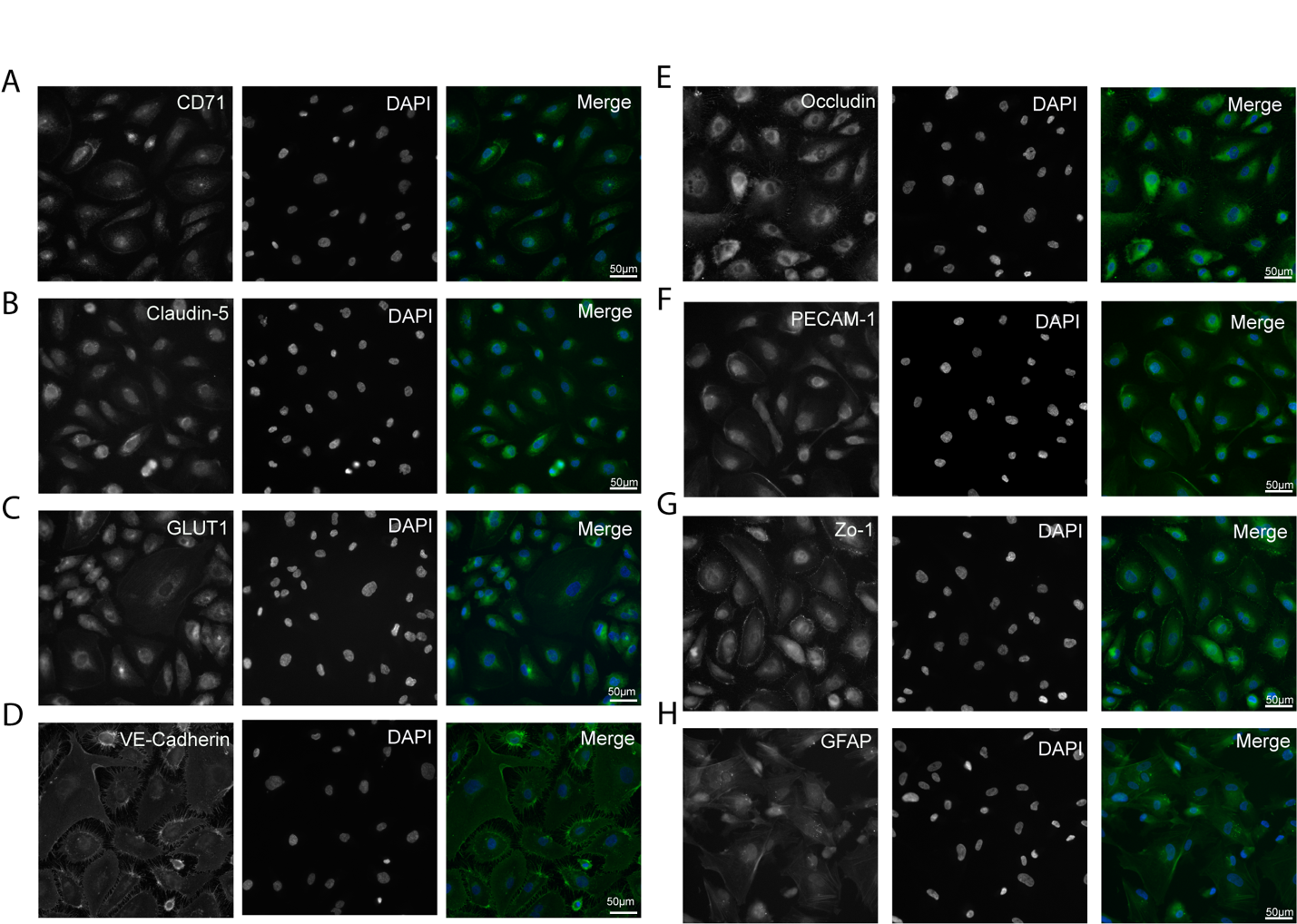

**Supplemental Figure 1. Brain Microvascular Endothelial Cells and Astrocytes Express Characteristic Markers**. Immunofluorescent microscopy was performed to evaluate expression of anticipated markers in (A-G) primary human brain microvascular endothelial cells and (H) primary human astrocytes. (Left panels) Antibodies with specificity to (A) CD71, (B) claudin-5, (C) GLUT-1, (D) VE-Cadherin, (E) occludin, (F) PECAM-1, (G) Zo-1, and (H) GFAP were coupled to Alexa Fluor 488 for analysis. (Middle panels) DAPI was used to visualize nucleus. (Right panels) Merge depicts the combined signal for proteins of interest (green) and DAPI (blue). Representative images, out of 20 independent images, are shown. All scale bars = 50 μm.

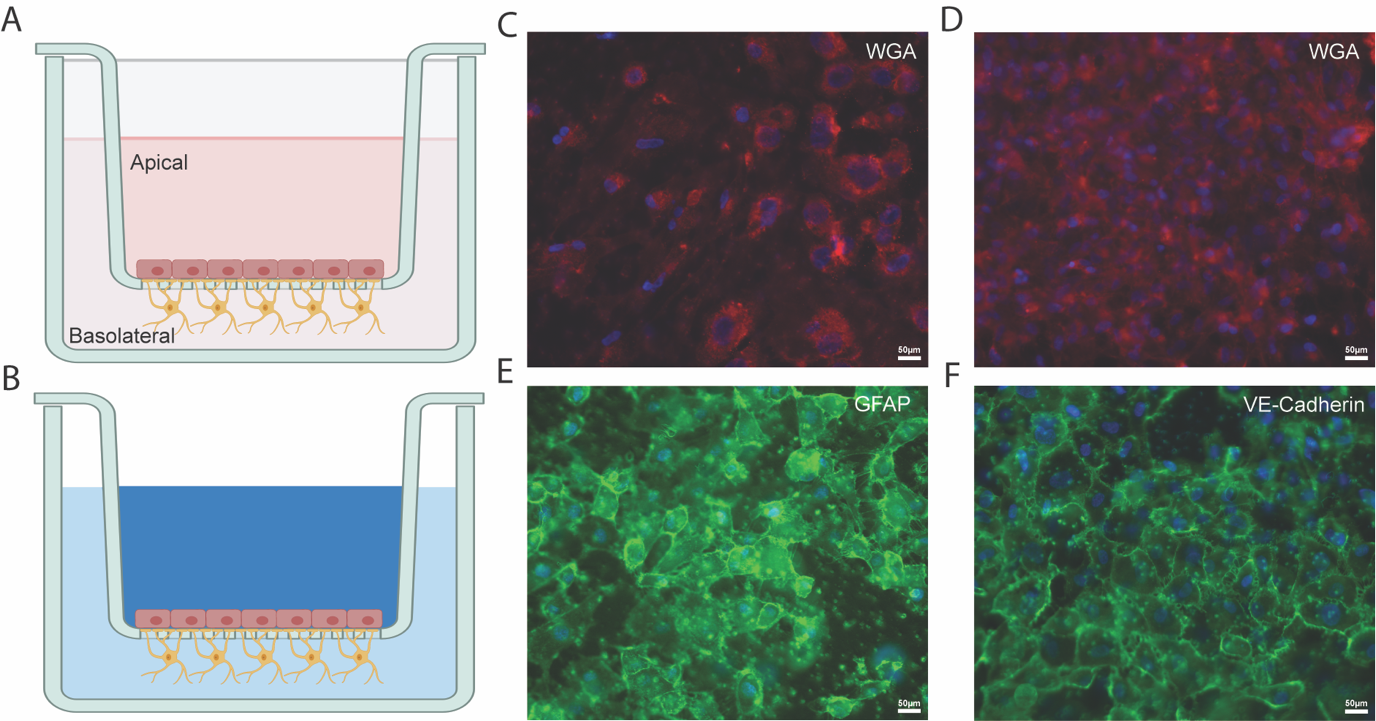

**Supplemental Figure 2. *In vitro* Model of the Human BBB.** (A) Schematic representation of our transwell BBB model where primary human brain microvascular endothelial cells are seeded on the upper, apical compartment and primary human astrocytes are seeded on the underside of a polycarbonate membrane with 3 μm portes in the basolateral compartment. (B) Schematic representation of albumin permeability assay, where EBA dye is added to the apical portion and permitted to pass to the basolateral side for 30 minutes at 37°C, 5% CO_2_. The media in the basolateral side is collected and spectrophometrically read at OD_620_ to evaluate BBB permeability. (C-F) The polycarbonate membrane from the BBB model was collected, immunostained, and immunofluorescent microscopy performed. (C-D) Wheat germ agglutinin (WGA) depicts cell morphology in red and (E-F) demonstrates expression of the astrocyte and endothelial cell markers GFAP and VE-Cadherin, respectively in green. DAPI was used to visualize nucleus (blue). Representative images, out of 3-5 independent images, are shown. All scale bars = 50 μm.

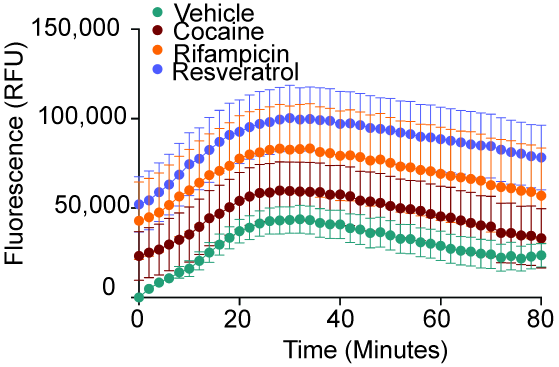

**Supplemental Figure 3. Complete Duration of CYP3A4 Metabolic Activity Assay**. Endothelial cells were pre-treated with cocaine (10 μM, burgundy), rifampicin (1 μM, yellow), resveratrol (10 μM, lavender), or vehicle (teal) for 24 hours, after which time the cells were loaded with BFC (2 μM). The enzymatic capacity of CYP3A4 to convert BFC to HFC was determined for 80 minutes as determined by fluorometric quantitation at excitation and emission wavelengths of 405/535 nm. Twelve independent experiments that contained eight technical replicates per condition were performed. Data are represented as mean ± standard deviation.

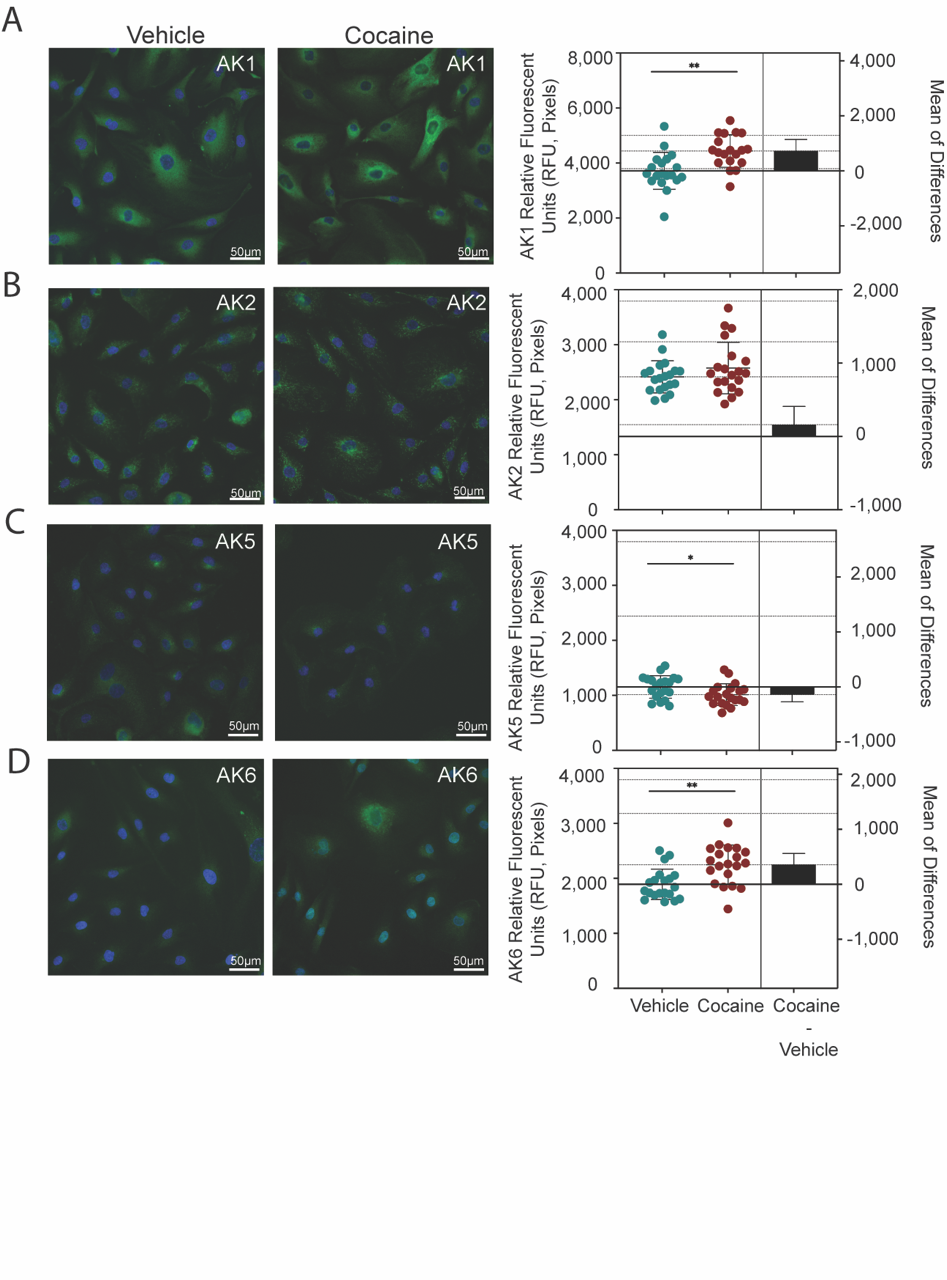

**Supplemental Figure 4. Cocaine Modulates AK1, AK5, and AK6 Expression**. Immunofluorescent microscopy was performed to evaluate (A) AK1, (B) AK2, (C) AK5, and (G) AK6 (green) following treatment with cocaine (10 μM, right) or vehicle (left) for 24 hours. DAPI was used to visualize nucleus (blue). One paired representative image, out of 20 individual images, are shown. All scale bars = 50 μm. Quantification of the fluorescent signal from immunofluorescent microscopy was performed for endothelial cells treated with cocaine (10 μM, burgundy) or vehicle (teal) for 24 hours. Twenty independent experiments (represented by individual dots) were performed. Estimation plots are shown where the left y-axis denotes relative fluorescent intensity (RFU, pixels) and the right y-axis reflects the effect size (black bar), which is the difference between means of each condition. Data are represented as mean ± standard deviation. *p<0.05. **p<0.01. Unpaired T-test was performed.
